# Pancreatic cancer patient-derived organoids capture therapy response and tumor evolution

**DOI:** 10.1101/2025.04.24.650216

**Authors:** Johann Gout, Lukas Perkhofer, Jessica Lindenmayer, Yazid J. Resheq, Julian D. Schwab, Johann M. Kraus, Elodie Roger, Dharini Srinivasan, Eleni Zimmer, Benjamin Mayer, Ralf Marienfeld, Nadine T. Gaisa, Julia K. Kühlwein, Mark Hänle, Thomas J. Ettrich, Karin M. Danzer, Alica K. Beutel, Bo Kong, Christoph Michalski, André L. Mihaljevic, Nuh N. Rahbari, Marko Kornmann, Hans A. Kestler, Thomas Seufferlein, Alexander Kleger

## Abstract

Patient-derived organoids (PDOs) reflect parental tumor features and may represent promising avatars for prognosticating drug response. Here, we recruited 169 patients with pancreatic cancer (PC) and established a living biobank including 83 pharmacotyped PDOs isolated from primary and metastatic, treatment-naïve and pretreated PCs. In a core facility setting, the pharmacotyping success rate was 61.5%, with an unmet turnaround time of 32 days. Forty-six patients who underwent a total of 94 therapeutic lines were analyzed, resulting in a pharmacotyping-patient response matching rate of 73.4%. Sensitivity, specificity, positive and negative predictive values were 85.0%, 64.8%, 64.2%, and 85.4%, respectively. Tracing clonal evolution in longitudinal biopsies uncovered therapy-induced genetic alterations and single-nucleus multiomics identified transcriptomic and epigenetic changes associated with abnormal FGF signaling during treatment in one particular tracked study case. Our findings highlight the potential of PDOs as robust tools for drug response prediction and patient modeling to advance functional precision medicine.

## INTRODUCTION

Pancreatic adenocarcinoma (PC) remains a significant clinical challenge, characterized by a bleak 5-year survival that has only recently reached 13%,^1,2^ underscoring the urgent need for innovative and effective therapeutic strategies. The poor prognosis is largely due to late-stage diagnoses, resulting from the lack of specific early symptoms, combined with the aggressive tumor biology, and considerable inter and intratumor heterogeneity.^3,4^ Although temporarily effective, combinatorial cytotoxic regimens are associated with toxicity impacting quality of life, and secondary resistance hindering longer-term treatment. Additionally, primary resistance to untargeted therapies underscores the critical need for reliable predictive models or biomarkers to guide therapeutic decision making. Despite up to 25% of pancreatic cancer patients harboring actionable mutations, practical therapeutic implementation is lagging due to various factors, resulting in less than 2% receiving matched therapies based on genomic profiling.^5^ Consequently, conventional chemotherapy remains the primary PC treatment modality, with three standard-of-care first-line regimens applied largely based on the patient’s physical performance: gemcitabine plus nab-paclitaxel (GnP), NALIRIFOX (nanoliposomal irinotecan, 5-fluorouracil, leucovorin, and oxaliplatin), and (modified) FOLFIRINOX (folinic acid, 5-fluorouracil, irinotecan, and oxaliplatin).^6–10^ Further challenges arise as only around 50% of those patients treated with first-line therapy proceed to second-line treatment.^11^ The only approved targeted therapy for PDAC is olaparib, a DNA-damaging drug that acts synthetically lethal with *BRCA1/2* mutations, which occur in approximately 2-3% of cases.^7,12^ Additionally, advanced clinical trial data for mutant KRAS inhibitors is eagerly awaited, as promising (pre)clinical results continue to emerge.^13,14^ In summary, limited success of targeted therapies in real world underpins the complex and difficult-to-treat nature of PC and calls for functional precision oncology using patient-derived assays to bridge genotype and phenotype.^5,15^ Patient-derived xenograft (PDX) and patient-derived organoid (PDO) models have emerged as powerful tools in both basic and translational research.^15,16^ In this context, we have previously performed a head-to-head comparison of PDX and PDO responsiveness to the individual components of the two main chemotherapy regimens, revealing overall high concordance.^17^ While PDX better replicate the intricate tumor microenvironment, their time-consuming and labor-intensive nature limits their suitability for timely preclinical screening. In contrast, PDOs offer a practical method for *in vitro* pharmacotyping within a reasonable timeframe, holding promise for overcoming the limitations posed by PDX models.^17,18^ PDO-derived gene signatures have shown predictive value in chemotherapy sensitivity, and *in vitro* drug responses align with various clinical outcomes in PC and educated tumor vaccine development.^19–23^ We recently conducted a small prospective feasibility trial for real-time isolation of PDOs, which revealed diverse responses to standard-of-care drugs. We classified the organoid lines as high, intermediate, or low responders and developed a prediction model that accurately forecasted responses to both first-line and second-line treatments in treatment-naïve patients.^18,21,24^ Furthermore, a recent PDO-based prospective trial correctly identified putative chemotherapy hits for 91% of patients.^25^ However, accuracy decreased in pretreated patients, especially for those with multiple prior chemotherapy lines, but sample size remained low to draw final conclusions. Still, several questions remain unanswered: (i) how feasible are quality improvement initiatives, such as centralized core facilities and standardized procedures in real-world? Is it possible (ii) to deliver pharmacotyping data within a clinically meaningful time frame and (iii) to simultaneously extend drug panels, to assist treatment decision-making beyond standards of care and truly foster precision oncology? (iv) How do PDO pharmacotyping profiles correlate with patient responses in a large real-world cohort of PC patients? (v) Is it possible to capture therapy-driven tumor evolution through longitudinal sampling of PDOs? (vi) Can the cellular heterogeneity of pancreatic cancer and its associated molecular pathways be elucidated through epigenomic and gene expression profiling of PDOs, potentially revealing previously hidden insights? We broadened the scope of our trial by assessing the potential of PDOs as a diagnostic tool to predict drug responses in routine clinical settings. The current study, conducted within the frame of an organoid core facility, represents the largest prospective PC PDO cohort to date, incorporating multiple subsequent lines of treatment from pharmacotyped patients, demonstrating an as yet unmet turnaround time with high accuracy. Additionally, we introduce a novel dimension to our study by integrating a comprehensive analysis of clonality through whole exome and single-nucleus multiome sequencing. This approach sheds light on the intricacies of tumor evolution and response dynamics at bulk and single-cell level, respectively. Our multifaceted and standardized methodology provides deeper insights into the potential of PDOs for informing precision medicine in PC.

## RESULTS

### Study design and patient characteristics

A PC organoid registry was established in 2019,^24^ initially sampling specimens for organoid derivation during an exploratory phase. This was succeeded by the implementation of a centralized core facility-based structure. We enrolled 169 patients and successfully established a total of 87 PDO lines (**Figure 1A**). Baseline demographics and characteristics of all enrolled patients with at least one successfully derived organoid line are summarized in **Table 1**. Diagnostic pathology confirmed the presence of exocrine pancreatic malignancy in all samples included in the study. Among the 78 patients from whom a PDO line was established – including eight longitudinal patients with multiple organoids (nine additional PDOs for a total of 87 PDOs) – 75 were diagnosed with pancreatic ductal adenocarcinoma. The remaining three patients were diagnosed with rare subtypes of pancreatic tumors, which included one case of carcinoma derived from a solid pseudopapillary neoplasm of the pancreas, one squamous cell carcinoma, and one undifferentiated anaplastic carcinoma (**Table 1**). Most of the PC were staged as advanced diseases: 4 stage I (A+B; median overall survival (mOS): undefined), 11 stage II (A+B; mOS: 383.0 days), 1 stage III (mOS: undefined), and 62 stage IV (mOS: 249.0 days) (**Figure S1A**). Biological samples for organoid derivation were collected from either the primary pancreatic tumor (n=30) or metastases (n=48) (**Table 1**). Metastasis-derived organoid lines were obtained from liver (n=43), peritoneum (n=3), and lymph node metastatic sites (n=2). Notably, 25 biological samples (14.8%) had to be excluded due to excessive fibrotic tissue, bacterial contamination, or heavy erythrocyte contamination. Fifty-seven PC patients were sampled at the time of the primary diagnosis (treatment-naïve), while 21 patients were biopsied at tumor progression stage (pretreated). Among the pretreated patients at the time of biopsy, seven had received one prior systemic therapy, and 14 were already heavily pretreated (≥2 lines). Therapy response data were available for 46 patients, who underwent a total of 94 therapy lines, enabling organoid-based predictions across all treatment settings: neoadjuvant (n=5), adjuvant (n=7), and palliative (n=82) (**Figure 1A**).

**Figure 1.**
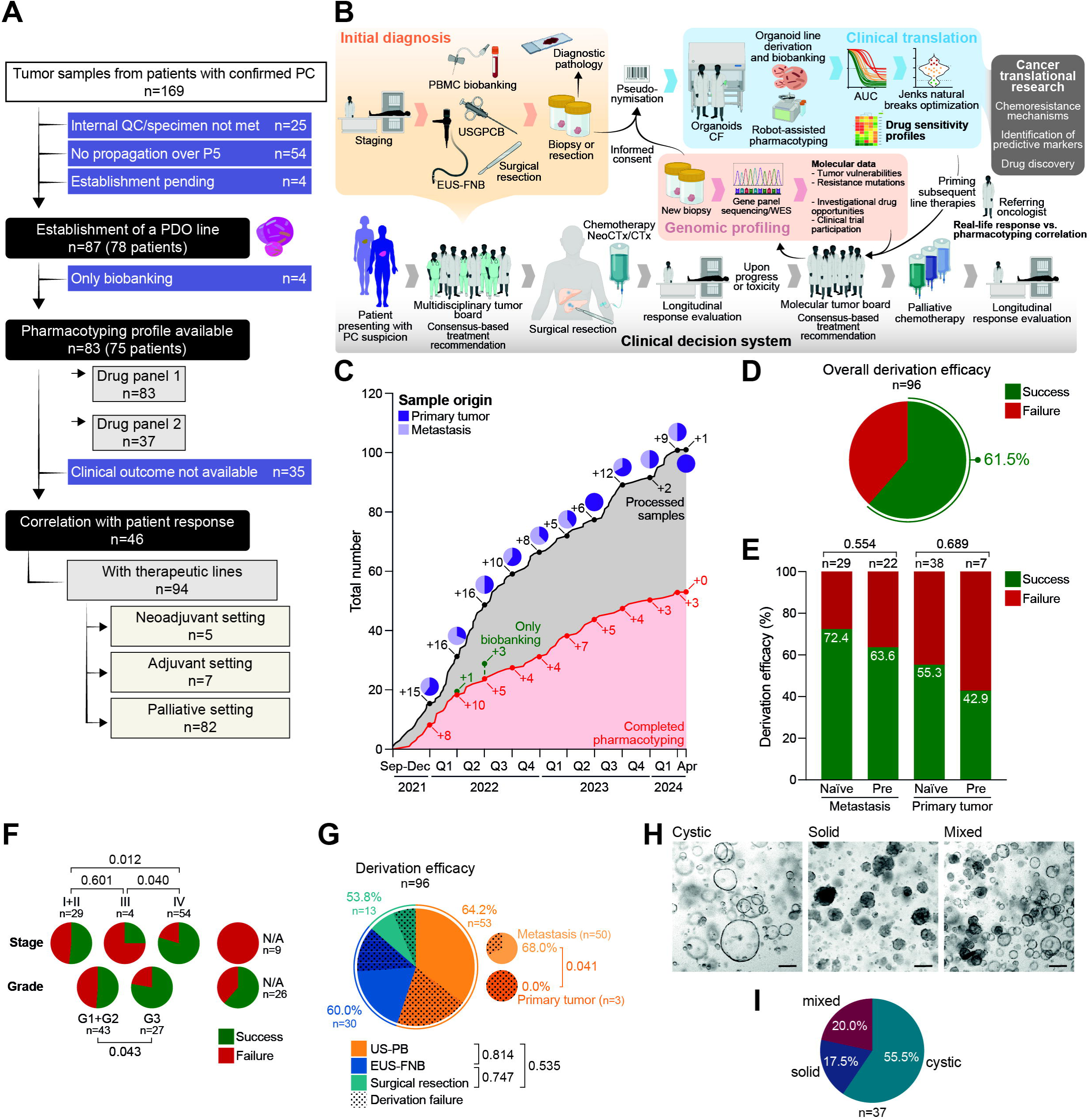
Workflow and performance of an organoid core facility operating in translational pancreatic cancer research. (A) Schematic representation of the study workflow. (B) Flow diagram of the study. (C) Organoid core facility pancreatic cancer-related workload history over the course of the 32 months following its creation. (D–G) Pie charts depicting the overall PDO line derivation efficacy (D) and the corresponding derivation efficacies according to biological material origin and treatment status (E), to the tumor stage and grade (F), and to the sampling method (G). (H–I) Brightfield images of PDO lines morphology (H) and pie chart illustrating the distribution of the morphological characteristics in the organoid living biobank used in the study (I). Scale bars, 100 µm. AUC, area under the curve; CF, core facility; CTx, chemotherapy; EUS-FNB, endoscopic ultrasound-guided fine needle biopsy; N/A, not available; NeoCTx, neoadjuvant chemotherapy; ns, not significant; PBMC, peripheral blood mononuclear cells; PC, pancreatic cancer; PDO, patient-derived organoid; Pre, pretreated; Q, quarter; QC, quality control; US-PB, ultrasound-guided percutaneous biopsy; WES, whole exome sequencing.

**Table 1.**
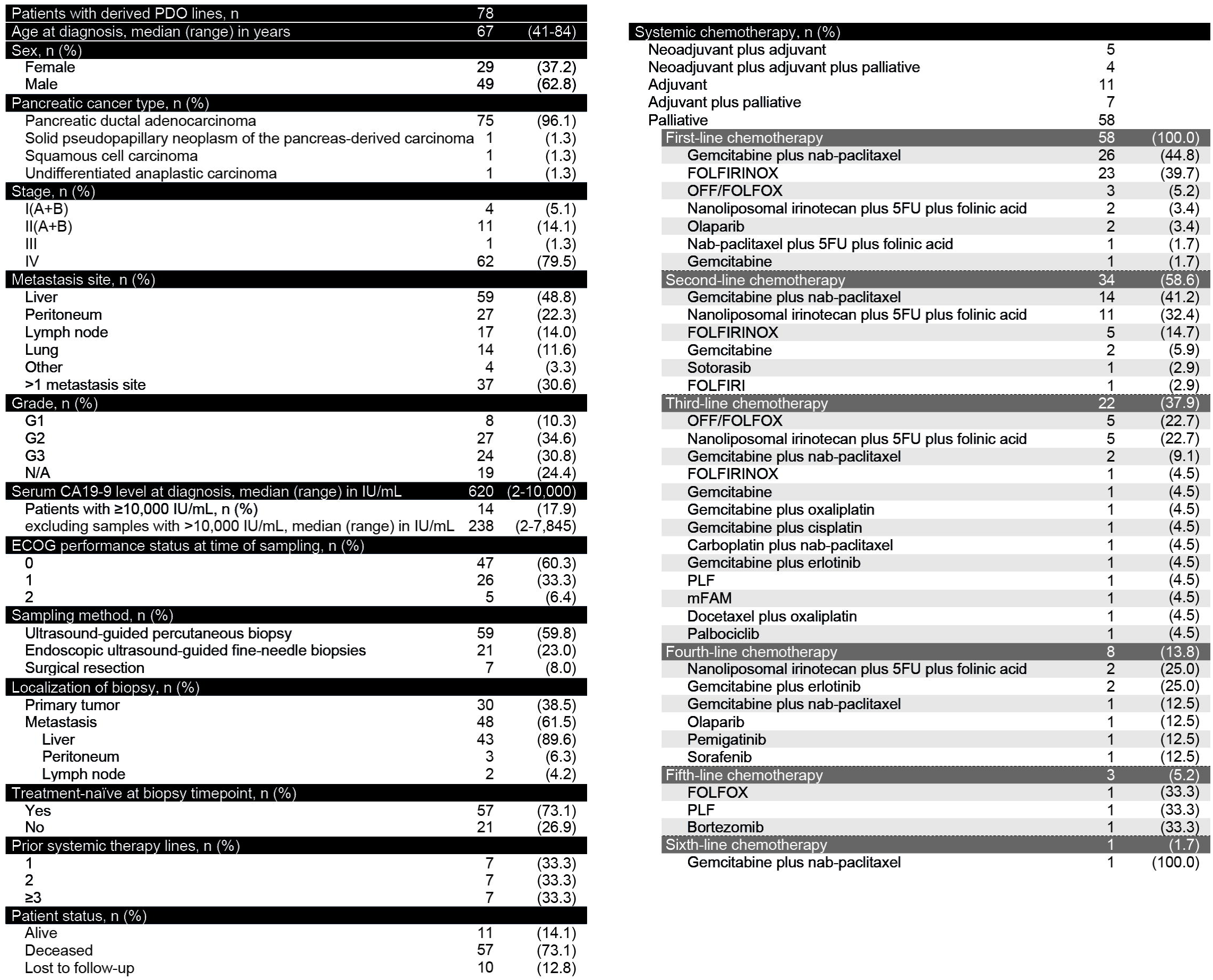
Patient characteristics. 5FU, 5-fluorouracil; ECOG, eastern cooperative oncology group; FOLFIRI, folinic acid, 5-fluorouracil, and irinotecan; FOLFIRINOX, folinic acid, 5-fluorouracil, irinotecan, and oxaliplatin; FOLFOX, folinic acid, 5-fluorouracil, and oxaliplatin; mFAM, 5-fluorouracil, doxorubicin, and mitomycin C; N/A, not available; nab-paclitaxel, nanoparticle albumin-bound paclitaxel; OFF, oxaliplatin, 5-fluorouracil, and folinic acid; PDO, patient-derived organoid; PLF, cisplatin, folinic acid, and 5-fluorouracil.

|  |  |  |
| --- | --- | --- |
| Patients with derived PDO lines, n | 78 |  |
| Age at diagnosis, median (range) in years | 67 | (41-84) |
| Sex, n (%) |  |  |
| Female | 29 | (37.2) |
| Male | 49 | (62.8) |
| Pancreatic cancer type, n (%) |  |  |
| Pancreatic ductal adenocarcinoma | 75 | (96.1) |
| Solid pseudopapillary neoplasm of the pancreas-derived carcinoma | 1 | (1.3) |
| Squamous cell carcinoma | 1 | (1.3) |
| Undifferentiated anaplastic carcinoma | 1 | (1.3) |
| Stage, n (%) |  |  |
| I(A+B) | 4 | (5.1) |
| II(A+B) | 11 | (14.1) |
| III | 1 | (1.3) |
| IV | 62 | (79.5) |
| Metastasis site, n (%) |  |  |
| Liver | 59 | (48.8) |
| Peritoneum | 27 | (22.3) |
| Lymph node | 17 | (14.0) |
| Lung | 14 | (11.6) |
| Other | 4 | (3.3) |
| >1 metastasis site | 37 | (30.6) |
| Grade, n (%) |  |  |
| G1 | 8 | (10.3) |
| G2 | 27 | (34.6) |
| G3 | 24 | (30.8) |
| N/A | 19 | (24.4) |
| Serum CA19-9 level at diagnosis, median (range) in IU/mL | 620 | (2-10,000) |
| Patients with $\geq 10,000$ IU/mL, n (%) | 14 | (17.9) |
| excluding samples with $>10,000$ IU/mL, median (range) in IU/mL | 238 | (2-7,845) |
| ECOG performance status at time of sampling, n (%) |  |  |
| 0 | 47 | (60.3) |
| 1 | 26 | (33.3) |
| 2 | 5 | (6.4) |
| Sampling method, n (%) |  |  |
| Ultrasound-guided percutaneous biopsy | 59 | (59.8) |
| Endoscopic ultrasound-guided fine-needle biopsies | 21 | (23.0) |
| Surgical resection | 7 | (8.0) |
| Localization of biopsy, n (%) |  |  |
| Primary tumor | 30 | (38.5) |
| Metastasis | 48 | (61.5) |
| Liver | 43 | (89.6) |
| Peritoneum | 3 | (6.3) |
| Lymph node | 2 | (4.2) |
| Treatment-naïve at biopsy timepoint, n (%) |  |  |
| Yes | 57 | (73.1) |
| No | 21 | (26.9) |
| Prior systemic therapy lines, n (%) |  |  |
| 1 | 7 | (33.3) |
| 2 | 7 | (33.3) |
| $\geq 3$ | 7 | (33.3) |
| Patient status, n (%) |  |  |
| Alive | 11 | (14.1) |
| Deceased | 57 | (73.1) |
| Lost to follow-up | 10 | (12.8) |
| Systemic chemotherapy, n (%) |  |  |
| Neoadjuvant plus adjuvant | 5 |  |
| Neoadjuvant plus adjuvant plus palliative | 4 |  |
| Adjuvant | 11 |  |
| Adjuvant plus palliative | 7 |  |
| Palliative | 58 |  |
| First-line chemotherapy | 58 | (100.0) |
| Gemcitabine plus nab-paclitaxel | 26 | (44.8) |
| FOLFIRINOX | 23 | (39.7) |
| OFF/FOLFOX | 3 | (5.2) |
| Nanoliposomal irinotecan plus 5FU plus folinic acid | 2 | (3.4) |
| Olaparib | 2 | (3.4) |
| Nab-paclitaxel plus 5FU plus folinic acid | 1 | (1.7) |
| Gemcitabine | 1 | (1.7) |
| Second-line chemotherapy | 34 | (58.6) |
| Gemcitabine plus nab-paclitaxel | 14 | (41.2) |
| Nanoliposomal irinotecan plus 5FU plus folinic acid | 11 | (32.4) |
| FOLFIRINOX | 5 | (14.7) |
| Gemcitabine | 2 | (5.9) |
| Sotorasib | 1 | (2.9) |
| FOLFIRI | 1 | (2.9) |
| Third-line chemotherapy | 22 | (37.9) |
| OFF/FOLFOX | 5 | (22.7) |
| Nanoliposomal irinotecan plus 5FU plus folinic acid | 5 | (22.7) |
| Gemcitabine plus nab-paclitaxel | 2 | (9.1) |
| FOLFIRINOX | 1 | (4.5) |
| Gemcitabine | 1 | (4.5) |
| Gemcitabine plus oxaliplatin | 1 | (4.5) |
| Gemcitabine plus cisplatin | 1 | (4.5) |
| Carboplatin plus nab-paclitaxel | 1 | (4.5) |
| Gemcitabine plus erlotinib | 1 | (4.5) |
| PLF | 1 | (4.5) |
| mFAM | 1 | (4.5) |
| Docetaxel plus oxaliplatin | 1 | (4.5) |
| Palbociclib | 1 | (4.5) |
| Fourth-line chemotherapy | 8 | (13.8) |
| Nanoliposomal irinotecan plus 5FU plus folinic acid | 2 | (25.0) |
| Gemcitabine plus erlotinib | 2 | (25.0) |
| Gemcitabine plus nab-paclitaxel | 1 | (12.5) |
| Olaparib | 1 | (12.5) |
| Pemigatinib | 1 | (12.5) |
| Sorafenib | 1 | (12.5) |
| Fifth-line chemotherapy | 3 | (5.2) |
| FOLFOX | 1 | (33.3) |
| PLF | 1 | (33.3) |
| Bortezomib | 1 | (33.3) |
| Sixth-line chemotherapy | 1 | (1.7) |
| Gemcitabine plus nab-paclitaxel | 1 | (100.0) |

### Foundation and performance of an organoid core facility

In September 2021, we launched the Core Facility (CF) Organoids at Ulm University to standardize processes with comprehensive standard operating procedures within a centralized infrastructure (**Document S1**). Here, the digitally preregistered tissue sampling is conducted from primary or metastatic sites during surgical resection or through endoscopic or percutaneous ultrasound-guided biopsy. Biospecimens are transported to the CF, pseudonymized, and then processed for PDO derivation, which contributes to the establishment of a living biobank. For PC pharmacotyping, two drug panels are available: panel 1, which includes standard-of-care (SOC) drugs (gemcitabine, paclitaxel, irinotecan, 5-fluorouracil, oxaliplatin) and panel 2, which tests additional targeted therapies (olaparib, erlotinib, palbociclib) and cisplatin. However, sensitivity to therapeutic agents is not yet being used to guide clinical decision. Targeted molecular testing, including genomic profiling, is conducted in accordance with the recommendations from the molecular tumor board (MTB) to identify potential therapy-relevant mutations and signaling pathways. The insights gained from this testing are integrated into consensus-based treatment decision-making within the MTB, primarily focusing on therapies beyond the first line. This workflow seamlessly combines diagnostics, organoid technology, and personalized medicine, ultimately aiming to enhance treatment outcomes for pancreatic cancer patients (**Figure 1B**). The CF collected and processed 96 PC samples for organoid derivation. Among these, 59 PDO lines were successfully established, propagated, and biobanked, resulting in an overall derivation success rate of 61.5% (**Figures 1C and 1D**). Organoid generation tended to be more efficient in biopsies from metastatic lesions than from primary tumor (**Figures 1E and S1B**), with the highest success rate in liver metastasis. However, possible disparities between distinct metastatic sites could not be entirely ruled out due to limited sample sizes (**Figure S1B**). Interestingly, pretreatment status did not significantly affect the derivation rate (**Figures 1E and S1C**). Furthermore, derivation efficacy showed no correlation with overall survival rates or with the time intervals between biopsy and either death or loss to follow-up (**Figures S1D and S1E**). Nevertheless, biological samples from metastatic and grade 3 cancers tended to have higher take rates (**Figure 1F**). Tumor cellularity and total tissue digestion to single-cell level are key limiting factors for organoid derivation.^26^ Therefore, we examined the efficacy of different sampling methods (**Figure 1G**). The highest PDO establishment rate was procured by ultrasound-guided percutaneous biopsies (US-PB), reflecting good results obtained with metastatic lesions (94.0% of samples) (**Figure S1B**). Surgical specimens exhibited the lowest success rate, likely due to their lower average tumor cell content.^24^ Endoscopic ultrasound-guided fine-needle biopsies (EUS-FNBs) performed moderately better in deriving stable organoid lines from primary tissues (**Figure 1G**). Noteworthy, all three primary tumor biopsies originating from an ultrasound-guided percutaneous puncture failed to produce an organoid line, demonstrating no superior performance of this technique (**Figure 1G**). PC organoid lines can vary significantly in morphology.^27,28^ We characterized 37 PDOs morphologically and categorized them into three patterns: cystic, solid, and mixed (**Figure 1H**). The most prevalent morphological subtype in our PC organoid living biobank was the cystic phenotype, followed by mixed and solid types (**Figure 1I**), showing a diverse distribution like previously described libraries.^19,29^

### Patient-derived organoids capture pharmacotypes in a clinically meaningful timeframe

To validate our CF-based workflow (**Figure 1B**) against the previously established manual profiling method, we performed a series of drug tests on six PDOs (**Figure 2A**). While we noted some variations in the sensitivity response clusters for single-agent tests (7 out of 13), the overall sensitivity profiles obtained from both the robot-assisted and manual methods remained consistent, resulting in comparable final prediction scores. This consistency underscores the robustness of our model. A rapid turnaround time is crucial for enabling informed clinical decision-making.^16^ The miniaturization and automation of the assay reduced the turnaround time for pharmacotyping from a median of 53.0 days (range 21-126 days)^24^ to 32.0 days (range 17-90 days), while simultaneously enhancing working capacity (**Figures 2B and 2C**). Drug tests were successfully conducted as early as 17 days after biopsy processing, demonstrating a significant improvement over the conventional manual method. Notably, the first 85th percentile of pharmacotypes was delivered within the previously established median turnaround time (**Figure 2B**). In this context, the quality of the initial biological material and the tumor cell content are critical factors to consider. As shown by the derivation rates (**Figure 1G**), there were no significant differences in data delivery times across the various sampling methods (**Figure S2A**). However, a trend emerged indicating that pharmacotyping may be completed more quickly for samples derived from high-grade and metastatic tumors (**Figure S2B**). To probe our organoid-based precision oncology approach, we compiled the individual times to neoadjuvant, adjuvant, and palliative treatment initiation, as well as the time to first restaging CT scan and progression upon first-line chemotherapy in the metastatic cases (**Figure S2C**). The 32-day pharmacotyping delivery time was shorter than the median time to the first restaging CT scan, occurring at 64.0 days after the start of systemic treatment (range: 26-80 days; p<0.001). Moreover, this delivery time was also quicker than the median time to progression following the first line of chemotherapy, which was 151.0 days (range: 46-827 days; p<0.001). Additionally, pharmacotyping was available prior to the median time of systemic treatment initiation in the adjuvant setting (median 59.0 days; range 50-67 days; p=0.013). Simultaneously, miniaturizing the assay expanded drug screening to include a secondary list of chemotherapeutic agents, including three targeted therapies. This advancement allows for accurate screening of treatment options beyond standard therapies within a similar timeframe (**Figures 2C–2E**) and enables adjustment of treatment based on actionable mutations. It further underscores the potential of organoid avatars as valuable diagnostic tools for detecting early therapy failure and monitoring disease progression in patients with advanced pancreatic cancer.

**Figure 2.**
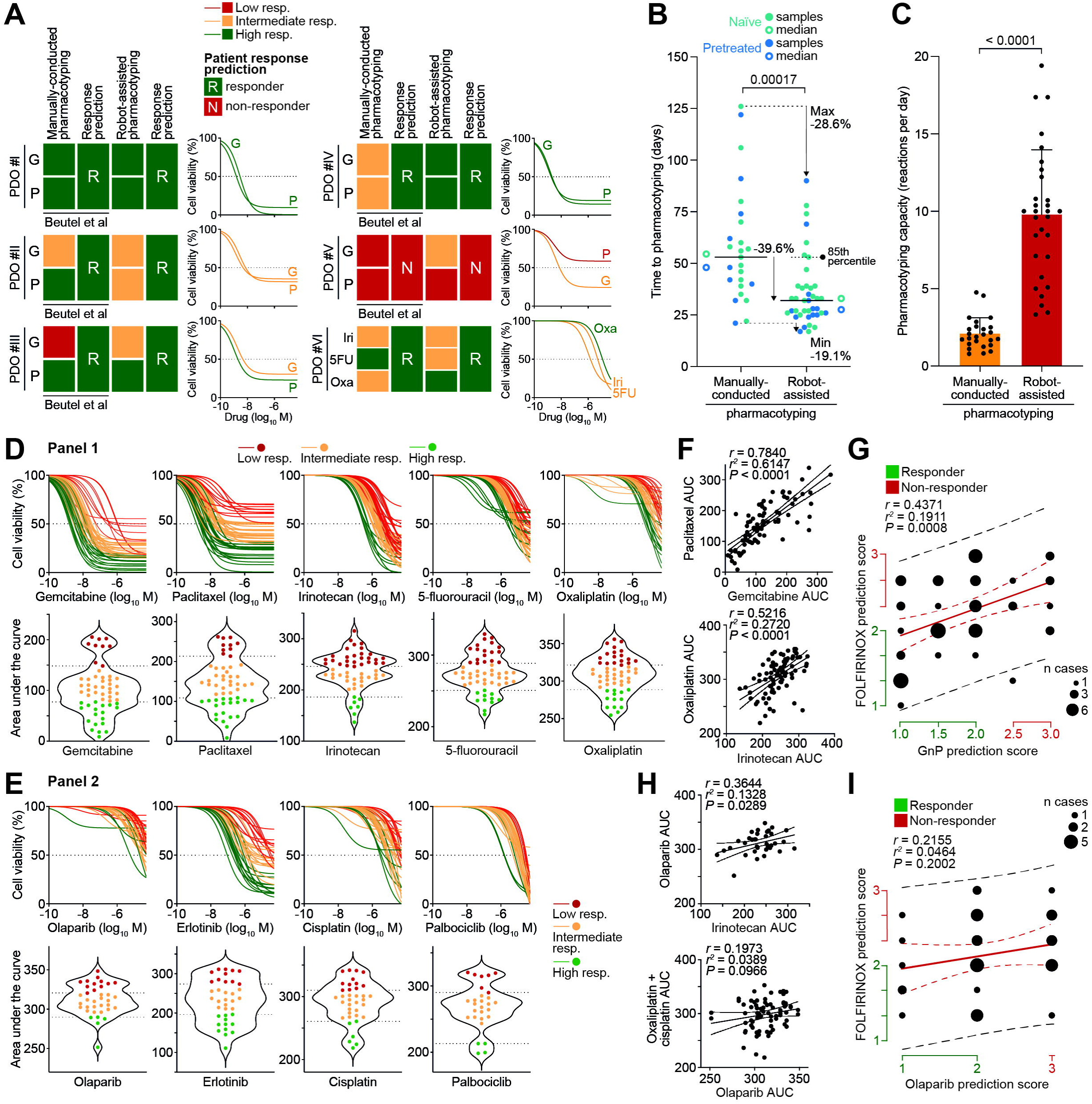
Standardized robot-assisted pharmacotyping workflow enhances feasibility and work capacity. (A) Prediction model robustness across both robot-assisted and manually-conducted procedures. (B–C) Time to organoid pharmacotyping (B) and work capacity (C) before and after standardization, miniaturization, and automation by an organoid core facility. (D) Dose-response curves (upper panels) and violin plots (lower panels) depicting corresponding areas under curves of gemcitabine, paclitaxel, irinotecan, 5-fluorouracil, and oxaliplatin (drug panel 1) in patient-derived organoid (PDO) lines. Dotted lines (lower panels) represent cut-off values determined by the Jenks natural breaks classification method. PDO lines were classified into three clusters: high responder (green), intermediate responder (orange), and low responder (red). (E) Dose-response curves (upper panels) and violin plots (lower panels) depicting corresponding areas under curves of olaparib, erlotinib, cisplatin, and palbociclib (drug panel 2) in PDO lines, as shown in (D). (F) Pearson correlation between individual organoid response to gemcitabine and paclitaxel (upper panel) and to oxaliplatin and irinotecan (lower panel). (G) Pearson correlation between pharmacotyping-based prediction scores for FOLFIRINOX and GnP in individual PDO lines. (H) Pearson correlation between individual organoid response to olaparib and irinotecan (upper panel) and to platins (oxaliplatin and cisplatin) and olaparib (lower panel). (I) Pearson correlation between pharmacotyping-based prediction scores for FOLFIRINOX and olaparib in individual PDO lines. 5FU, 5-fluorouracil; AUC, area under the curve; FOLFIRINOX, folinic acid, 5-fluorouracil, irinotecan, and oxaliplatin; G, gemcitabine; GnP, gemcitabine plus nanoparticle albumin-bound paclitaxel; Iri, irinotecan; Oxa, oxaliplatin; P, paclitaxel; resp., responder.

### Patient-derived organoids rationalize the use of standard-of-care regimens

Therapeutic profiling was conducted on 83 PDOs originating from 75 PC patients, including 15 longitudinal lines (**Figure 1A**). Drug response curves demonstrated significant heterogeneity in sensitivity to the nine individual compounds across all PDO lines. Our library of PDO lines effectively captured interpatient variations in therapy response, showcasing a wide range of chemosensitivity and resistance profiles (**Figures 2D and 2E**). We then analyzed the correlation between individual drug responses to provide an objective rationale for the clinical use of specific drug combinations.^8,30^ We found a robust positive correlation between the responses to gemcitabine and paclitaxel, as well as between oxaliplatin and irinotecan, reinforcing the rationale for their clinical combination (**Figure 2F**). Among all five standard therapeutic agents, most responses exhibited a general correlation, except for irinotecan versus 5-fluorouracil and paclitaxel versus oxaliplatin (**Figure S2D**). Notably, this resulted in a significant correlation between the prediction scores for FOLFIRINOX and the combination of gemcitabine plus nab-paclitaxel, indicating that organoids may exhibit a predictable sensitivity or resistance to both treatment regimens (**Figure 2G**). In contrast, no significant correlation was found between the two platinum agents (**Figure S2D**). We further explored the potential relationships between sensitivity to the personalized therapy, PARP inhibitor olaparib, and other DNA-damaging agents. Interestingly, a positive correlation was identified only between responses to olaparib and irinotecan (**Figures 2H and S2D**). Additionally, there was no correlation between the prediction scores for FOLFIRINOX and olaparib within our genomically-unstratified patient cohort (**Figure 2I**).

### Accurate organoid-based response prediction translates into clinical benefit

Next, we aimed to correlate pharmacotypes of the full PDO registry with the clinical patient outcomes. Clinical data were available for correlation in only 46 pancreatic cancer patients, highlighting the severity of the disease and frequent deterioration before the first restaging (**Figure 1B**). In the palliative setting, we analyzed 82 paired *in vitro*-*in vivo* responses to 13 regimens across 44 patients (**Figure 3A**). Assessment of the treatment response was conducted following RECIST criteria (response evaluation criteria in solid tumors v1.1), as the best and first available surrogate for the evaluation of palliative line efficacy.^21,24,29,31^ Overall, organoid sensitivities positively correlated with therapy responses in 73.2% of cases (60 of 82). Prior treatment status did not significantly impact the match ratio (naïve PDOs: 74.1%, 43 of 58; pretreated PDOs: 70.8%, 17 of 24; p=0.788). Furthermore, prediction rates for response to second next or subsequent therapies were not significantly altered: immediate next-line treatments showed a 69.8% match rate (30 of 43), second next-line treatments reached 70.8% (17 of 24), and subsequent lines an 86.7% concordance (13 of 15; p=0.529). This highlights the clinical relevance of our organoid platform, particularly in guiding second and subsequent palliative therapeutic lines. We also evaluated the diagnostic performance of PDOs in prognosticating therapeutic response in the neoadjuvant setting (**Figure 3B**). Five treatment-naïve patients – three with borderline resectable tumors and two locally advanced tumors – were included in our study. Three outcome parameters were used to evaluate the *in vivo* pathological response to the treatment: serum CA19-9 levels, histopathology-based tumor regression grading, and resection margin. In four cases, we observed concordance between organoid and real-world drug responses, suggesting that PDO pharmacotyping may also be valuable for tailoring neoadjuvant treatment approaches. Our study also included six resected patients receiving adjuvant therapy (**Figure 3C**). Among the three cases that were previously part of the neoadjuvant cohort, only one yielded an organoid line derived post-resection. Due to feasibility considerations, we determined six-month disease-free survival the earliest clinical outcome parameter to link pharmacotyping data directly to responses to adjuvant therapy, categorizing recurrences that occur within the first six months as early.^32^ In this context, our organoid model achieved an overall prediction accuracy of 71.4% (5 of 7). Notably, of the two sequential PDOs established for patient #40, only the second pretreated line (#40.2), collected during surgery, showed resistance to the combination of gemcitabine and 5-fluorouracil (next line). In conclusion, PC patients in the adjuvant setting may benefit from organoid-guided treatment recommendations.

**Figure 3.**
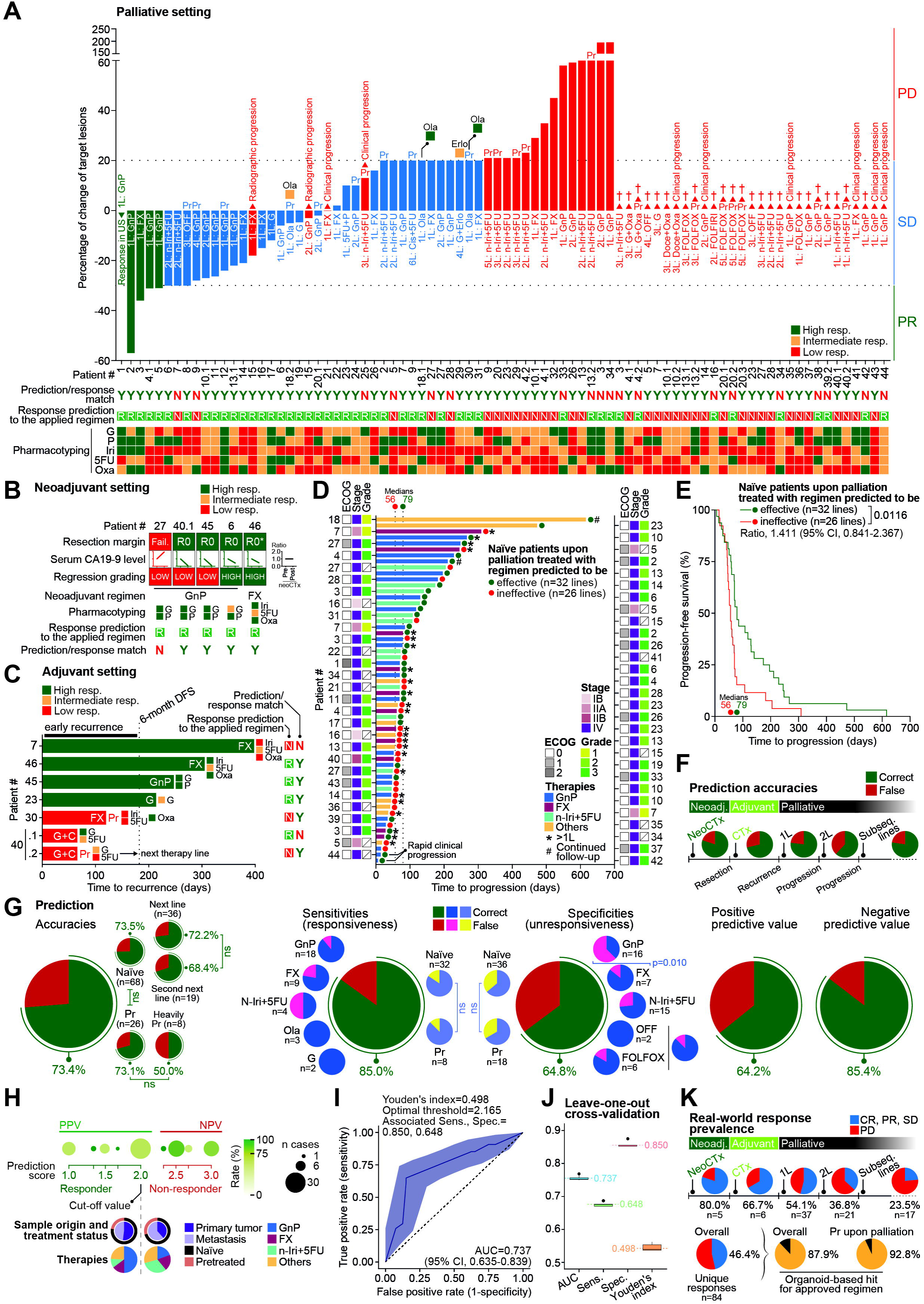
Patient-derived organoids, preclinical avatars to predict patient therapeutic response in pancreatic cancer. (A–C) Organoid pharmacotype and patient response correlation in the palliative (A), neoadjuvant (B), and adjuvant settings (C). R0*, margin-negative resection with one position of uncertain perineural infiltration. (D–E) Swimmer plot (D) and Kaplan-Meier analysis (E) showing progression-free survival of treatment-naïve PC patients upon palliation who received a regimen predicted to be effective or ineffective by our PDO pharmacotyping-based model. (F) Prediction accuracies per disease situation. (G) Performance metrics of our organoid core facility and prediction model. (H) Positive and negative predictive values per prediction score. (I–J) Receiver operating characteristic curve (I) and leave-one-out cross-validation (J) of our patient response classifier performance. (K) Tumor response prevalence per disease situation and hit identification rate for an approved combinatorial regimen. 1L, palliative first-line chemotherapy; 2L, palliative second-line chemotherapy; 3L, palliative third-line chemotherapy; 4L, palliative fourth-line chemotherapy; 5FU, 5-fluorouracil; 5L, palliative fifth-line chemotherapy; 6L, palliative sixth-line chemotherapy; AUC, area under the curve; C, capecitabine; CR, complete response; CTx, chemotherapy; DFS, disease-free survival; Doce, docetaxel; ECOG, eastern cooperative oncology group; Erlo, erlotinib; FOLFIRI, folinic acid, 5-fluorouracil, and irinotecan; FOLFOX, folinic acid, 5-fluorouracil, and oxaliplatin; FX, FOLFIRINOX (folinic acid, 5-fluorouracil, irinotecan, and oxaliplatin); G, gemcitabine; GnP, gemcitabine plus nanoparticle albumin-bound paclitaxel; Iri, irinotecan; n-Iri, nanoliposomal irinotecan; Neoadj., neoadjuvant; NeoCTx, neoadjuvant chemotherapy; NPV, negative prediction value; ns, not significant; OFF, oxaliplatin, 5-fluorouracil, and folinic acid; Ola, olaparib; Oxa, oxaliplatin; P, paclitaxel; PD, progressive disease; PDO, patient-derived organoid; PPV, positive prediction value; Pr, pretreated; PR, partial response; resp., responder; SD, stable disease; Sens., sensitivity; Spec., specificity; Subseq., subsequent; US, ultrasound.

To further assess the potential clinical value of our PDO predictions, we examined the progression-free survival (PFS) of treatment-naïve patients with advanced PC (**Figures 3D and 3E**). Remarkably, the administration of a predicted effective treatment led to significantly longer PFS (medians: 79 versus 56 days) in the palliative setting. To investigate the intra-patient relevance of our model, we additionally employed the commonly used PFS ratio.^33,34^ This metric compares the PFS achieved by a new systemic treatment (PFS2) to the PFS interval of the last prior therapy on which the patient has experienced progression (PFS1). Clinical benefit has been defined as a PFS ratio (PFS2/PFS1) >1.3.^35^ The predictive accuracy of our algorithm for therapeutic sequences was high, with 11 out of 15 efficacies correctly predicted (**Figure S2E**). This is further supported by the median PFS ratio of 1.300 (range: 0.070-9.520), with only 17.9% of sequences exceeding the 1.3 cut-off (**Figure S2F**), indicating that achieving meaningful clinical benefit beyond the first palliative line remains a challenge. Nevertheless, this paired analysis demonstrated that organoids may effectively guide the selection of second and subsequent treatment lines following disease progression. The analysis of the organoid model prediction ability across different disease states (resectable, advanced, and upon recurrence or progression) showed no significant decrease in accuracy, and thus, highlighted the robustness and value of our approach for various therapeutic scenarios (**Figure 3F**).

### Patient-derived organoids robustly predict therapeutic response

Considering all scenarios, the overall accuracy of our approach reached 73.4% (**Figure 3G**). Surprisingly, the prediction rates using naïve and pretreated PDOs were remarkably similar. However, accuracy decreased for heavily pretreated organoids, although not significantly. Naïve PDOs demonstrated comparable abilities to accurately predict treatment response for the current or immediate next line, as well as for the second next therapy line, in accordance with our previous findings.^24^ We further assessed the validity of our model by calculating four objective performance metrics (**Figure 3G**). Our approach demonstrated an overall better prediction of chemoresponsiveness (sensitivity) than unresponsiveness (specificity), achieving 85.0% and 64.8%, respectively. We also identified discrepancies in match ratios among different regimens for both outcomes. Sensitivity to the GnP regimen was predicted most accurately, followed by FOLFIRINOX and n-Iri+5FU. Notably, naïve and pretreated organoids exhibited similar prediction rates. Resistance to GnP was significantly mispredicted (6 of 16 versus 29 of 38 for all other regimens; p=0.010), which may help explaining the poorer performance in the unresponsiveness group. Importantly, no bias was detected based on PDO treatment status. Associated positive and negative predictive values were 64.2% and 85.4%, respectively (**Figure 3G**). As the precision of our prediction model was obviously affected by the number of false positives in the GnP predictions, we challenged the relevance of the threshold segregating non-response from response. However, a finer analysis of both predictive values per possible score revealed no evident flaw regarding the prediction model or relevant bias in the analyzed subgroup pinpointing intrinsic characteristics perturbing GnP prediction (**Figure 3H**).

Finally, we generated a receiver operating characteristic (ROC) curve to examine the method predictive potential in identifying clinical responders and non-responders, at each possible score threshold (**Figure 3I**). Overall, our analysis of 46 patients and 94 therapeutic lines achieved an area under the ROC curve of 0.737 (95% CI, 0.635-0.839), indicating that the model’s diagnostic accuracy for predicting therapy response falls within the acceptable range (>0.7^36^). Youden’s index was then employed to identify the prediction score cut-off point that maximizes the combination of sensitivity and specificity in an unbiased fashion.^37^ The computed threshold that best discriminated between responders and non-responders was 2.165, with a maximum Youden’s index of 0.498. Strikingly, this selected cut-point constitutes the closest-to-optimal possible classifying prediction score.^24^ Accordingly, the associated sensitivity, specificity, positive and negative predictive values remained unchanged (respectively, 0.850, 0.648, 0.642, and 0.854), as when using our previously defined cut-off point of 2.0. Data-driven determination of the optimal cut-off may lead to overly optimistic estimates of sensitivity and specificity metrics, especially with small sample sizes.^38^ To address this, we conducted an unbiased leave-one-out cross-validation which further confirmed our model validity (sensitivity 0.8500 (95% CI, 0.8488-0.8512), specificity 0.6481 (95% CI, 0.6467-0.6496), Youden’s index 0.4981 (95% CI, 0.4963-0.5000)) (**Figure 3J**). In conclusion, our organoid-based model displayed acceptable accuracy in prognosticating patient therapy response through *in vitro* pharmacoprofiling.

Across the entire cohort, the overall tumor response rate was 46.4% (39 out of 84 unique outcomes), with notable variations among different therapy settings: 80.0% (4 of 5) for neoadjuvant, 66.7% (4 of 6) for adjuvant, and 42.5% (31 of 73) for palliative treatments (p=0.179) (**Figure 3K**). Remarkably, our pharmacotyping approach identified a potential susceptibility to at least one state-of-the-art agent in 96.4% of the lines (80 of 83), and to an approved therapy combination in 87.9% (73 of 83) (**Figure 3K; Figure S2G**). The five most commonly used therapies accounted for 83.0% of the total predictions (**Figure S2H**). Only three PDOs exhibited resistance to all screened standard antitumor agents. Importantly, this vulnerability discovery rate was still 92.8% (26 of 28) among the pretreated advanced PC patients – a group known for its particularly low response rates^8,39^ (**Figure 3K**) – suggesting that our guidance could significantly enhance patient clinical outcomes. Collectively, these findings highlight the value of our approach in informing clinical decision-making across various scenarios involving pancreatic cancer.

### Genetic alterations captured in PDOs derived from sequential biopsies

Inter- and intratumoral heterogeneity, along with clonal evolution, are major determinants of PC and contribute to the establishment of chemoresistance under therapeutic selection pressure.^40,41^ However, the extent to which genetic heterogeneity and therapy-driven evolution can be accurately captured by PDOs is still unclear. We conducted whole exome sequencing (WES) ± panel sequencing on the longitudinal PDO lines from seven individuals with PC. Biological material was collected sequentially either due to disease progression (#4, #10, #13, #18, #20, #40) or because a second biopsy was required when histopathological diagnosis was uncertain (#39) (**Figures S3A– S3F**). Six PDO pairs (#4, #10, #13, #18, #20, #39) were derived from liver metastases, while the #40 tandem was obtained from the primary tumor (**Figure 4A**). To comprehensively address the mutational dynamics across these sequential PDO lines, we investigated the overall tumor mutation burden (TMB), single nucleotide variants (SNVs), and insertions-deletions (indels). When examining patient-matched organoid lines, a trend was observed toward a decrease in TMB (5 of 7) and SNVs (6 of 7), while indel levels remained relatively stable (**Figure 4B**) suggesting that SNVs are relevant events during tumor evolution.^42^ The analysis of the paired PDOs revealed varying frequencies of genetic alterations in the four most prevalent pancreatic cancer driver genes (*KRAS*, *TP53*, *CDKN2A*, and *SMAD4*).^43^ In the first PDO set (#X.1), *KRAS* alterations were found in 3 out of 7 lines (42.9%), *TP53* alterations in 4 out of 7 lines (57.1%), *CDKN2A* aberrations in 2 out of 7 lines (28.6%), and *SMAD4* losses in 3 out of 7 lines (42.9%) (**Figure 4A**). In the subsequent PDOs (#X.2), alteration rates increased for *KRAS* and *SMAD4* to 57.1% and 71.4% respectively. Losses in *CDKN2A* persisted unchanged in the second PDO. PDO #4.1 and #4.2 displayed acquired *KRAS* and *TP53* mutations, while the other organoid series either had no mutations or acquired a different type of mutation in one of these genes (*TP53* splice site versus missense in #10; *CDKN2A* frameshift versus nonsense in #13; *SMAD4* loss versus missense in #39) indicating changes in clonality. Although PDO #4.1 did not present any pathogenic mutations, it exhibited a pancreatic cancer-like pharmacotyping profile, showing sensitivity toward GnP while demonstrating resistance to the individual FOLFIRINOX components. This suggests the need for either additional assessment of structural aberrations or higher sequencing depth to capture most mutations (**Figure 4A**). Along this line, *ARID1A* (#10.1) and *KDM6A* (#13.1) mutations^44,45^ were absent in the respective second PDOs, while *SMAD4* loss was captured in the second biopsy of patient #13. Of note, the low *KRAS* mutation rate (usually >90%) in the entire cohort can be attributed to at least two atypical PC cases with alternative core drivers: *CTNNB1* mutations, typically found in solid pseudopapillary neoplasms of the pancreas (#18)^46^ and an *FGFR2::TPR* fusion (#20).^47^ Additionally, mutations in other genes known to contribute to tumor progression, such as *BRCA2*, *MSH6*, *RREB1*, and *NF1*,^43^ were dynamically identified with patterns of acquisition or loss (**Figure 4A**). Interestingly, four of seven PC patients included in our longitudinal cohort harbored actionable mutations (**Figure 4C; Figure S3**). Among them, three received a molecularly matched therapy, with two experiencing a prolonged response (#18 and #20). In contrast, only 14.3% of patients with available sequencing data were found to be carriers of actionable mutations (5 out of 35). Notably, the mutation rates for *KRAS* and *TP53* were 97.4% and 57.5%, respectively (**Figures S4A–S4B**). The implementation of molecularly matched therapy in patients with actionable mutations resulted in a significantly longer median overall survival compared to the unmatched cohort receiving standard-of-care treatments (p=0.027) (**Figure 4D**). PC patients whose tumors had no actionable alterations tended to exhibit a shorter median overall survival compared to those who received tailored therapy (p=0.199). However, there was no significant difference in overall survival when compared to the unmatched therapy group (p=0.595). These findings are consistent with previous observations reported in the *Know Your Tumor* registry trial, which investigated primary tissues and further licenses PDOs for functional phenotyping.^5^

**Figure 4.**
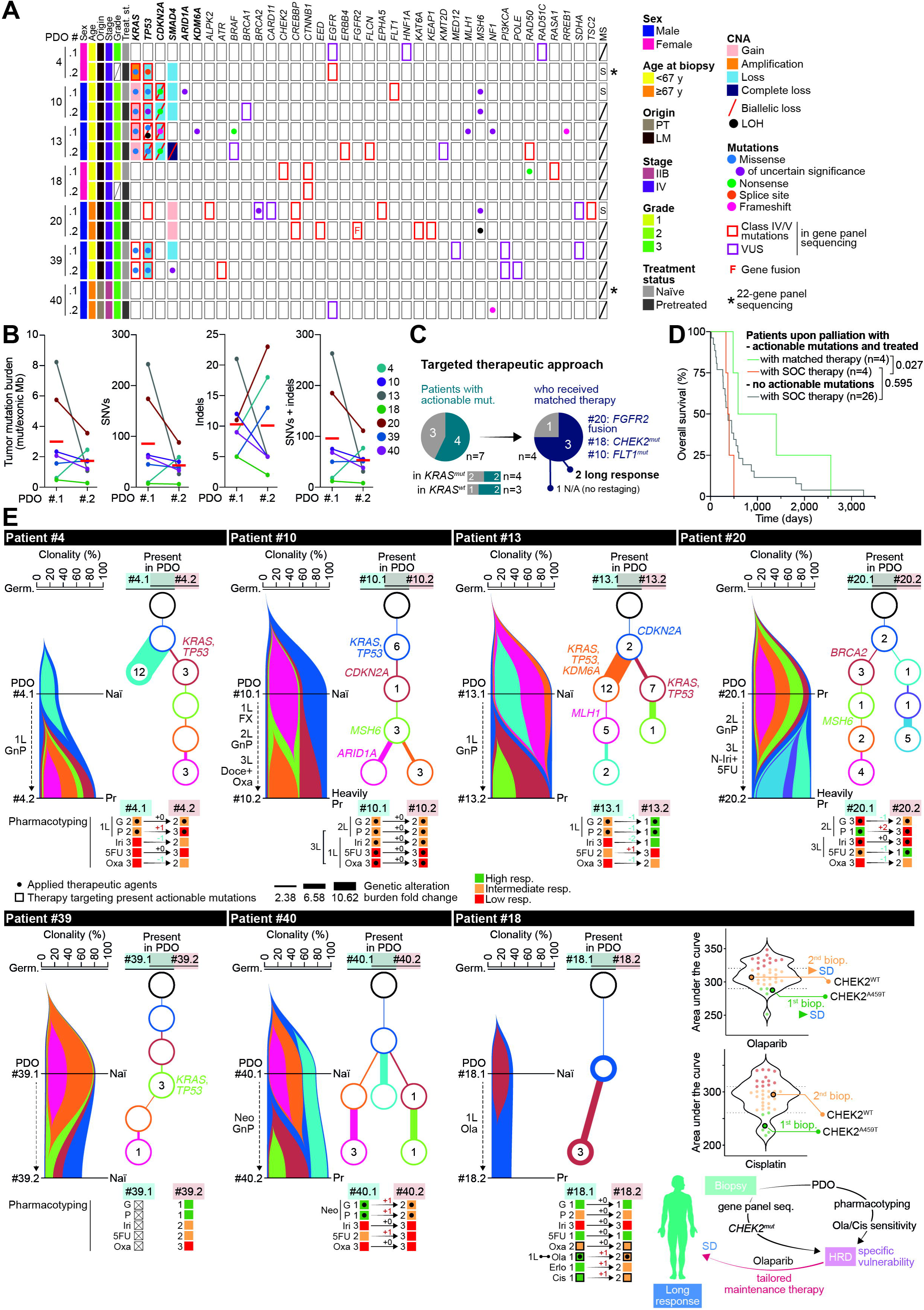
Patient-derived organoids capture tumor evolution. (A) Overview of the somatic mutations found in longitudinal patients by gene panel sequencing and whole exome sequencing. Cancer driver genes are highlighted in bold. (B) Tumor mutation burden in longitudinal PDO pairs. (C) Proportions of longitudinal patients whose tumors harbor actionable mutations and who received molecularly matched targeted therapy. (D) Kaplan-Meier analysis showing overall survival of PC patients whose tumors harbor or not actionable mutations, and who received a molecularly matched or a standard-of-care chemotherapy. (E) Clonal evolution of PDOs derived from longitudinal patients, shown as riverplots generated by SuperFreq analysis of whole exome sequencing. The x-axis represents the proportion of tumor cells in each subclone. The colored regions represent different subclones present in the longitudinal organoids. Evolutionary trees depicting the clonal situation in each PDO line are shown. Each node represents a subclone (corresponding to a different colored region). Branch thickness represents the total number of genetic alterations identified in each clone, as a fold change compared to its parental subclone. Representative cancer driver gene mutations harbored by each subclone are displayed. The total number of nonsynonymous mutations in each subclone are shown in the corresponding node. Single-agent pharmacotyping data obtained for longitudinal PDOs are shown with their corresponding sensitivity scores. For patient #18, individual PDO responses for olaparib and cisplatin are highlighted on the library violin plots, and a schematic representation depicts his particular situation. 1L, palliative first-line chemotherapy; 2L, palliative second-line chemotherapy; 3L, palliative third-line chemotherapy; 5FU, 5-fluorouracil; Cis, cisplatin; CNA, copy number alteration; Doce, docetaxel; FX, FOLFIRINOX (folinic acid, 5-fluorouracil, irinotecan, and oxaliplatin); G, gemcitabine; Germ., germline; GnP, gemcitabine plus nanoparticle albumin-bound paclitaxel; HRD, homologous recombination deficiency; Iri, irinotecan; LM, liver metastasis; LOH, loss of heterozygosity; MS, microsatellite; Mut., mutation; N/A, not available; Naï, treatment-naïve; Neo, neoadjuvant; n-Iri, nanoliposomal irinotecan; Ola, olaparib; Oxa, oxaliplatin; P, paclitaxel; PDO, patient-derived organoid; Pr, pretreated; PT, primary tumor; S, stable; SD, stable disease; SNV, single nucleotide variant; SOC, standard of care; TMB, tumor mutation burden; VUS, variant of unknown significance; wt, wild-type.

### Clonal evolution upon therapy and tumor progression

Assuming that these genetic alterations reflect the outgrowth of specific subclones with distinct mutational patterns, we compared the pharmacotyping profiles across these sequential PDO series for both standard-of-care and targeted treatments. Surprisingly, the observed shifts in genetic variations did not necessarily lead to a change in the pharmacotyping profile of the subsequently derived PDO line (**Figure 4E**). Specifically, PDO pairs #4, #20, #40, and #18 showed decreased sensitivity to at least one component of the treatment regimen, while PDO lines #10 and #13 displayed unusual patterns. These latter lines, derived longitudinally from two different metastatic lesions, exhibited a consistent pharmacotyping profile, suggesting either a lack of treatment-induced selective pressure or even an increased sensitivity to GnP, despite the patient having previously received this regimen (**Figure 4E**). This may suggest either the capture of a distinct subclone that is not critical to disease progression, or a limitation in interpretation stemming from the fact that patients often received multiple chemotherapies between the first and second biopsy.

To provide additional molecular context for these clinically-relevant changes, we monitored clonal evolution and heterogeneity in sequential PDOs by applying the SuperFreq algorithm^48^ (**Figure 4E**). All organoid lines displayed distinctly diverse patterns of clonality, ranging from 2 to 5 clones in the first PDO and from 1 to 5 clones in the subsequent PDO. This denotes a reduction in the degree of clonality observed in 4 of 7 organoid tandems upon disease course or therapy. In most cases, the emerging clones shared common alterations that formed the mutational backbone of the subsequently detected clones. The sole exception was the PDO series #4, where an increase in clonality was noted, coupled with the presence of different ancestral mutations in the second organoid line.

The changes in clonality were especially revealing in the sole patient who received targeted therapy. For patient #18, the detection of a pathogenic CHEK2^A459T^ mutation informed the decision to use the PARP1 inhibitor olaparib as a personalized maintenance therapy (**Figure S3F**). Simultaneously, PDO #18.1 was classified as highly sensitive to both olaparib and cisplatin, validating *in vitro* the patient’s vulnerability to DNA-damaging agents associated with the homologous recombination-related gene alteration (**Figure 4E**). Remarkably, the organoid-based prediction matched with the patient’s *in vivo* response. However, in the second biopsy taken during treatment, no CHEK2 alteration was found, and PDO #18.2 exhibited reduced sensitivity to both compounds. These changes in pharmacotyping were accompanied by the complete disappearance of an initially dominant clone, suggesting that olaparib may have eradicated particularly sensitive CHEK2-mutated cancer cells (**Figure 4E**).

### Longitudinal analysis of patient-derived organoids reveals key transcriptional and chromatin dynamics during therapy of a unique case

Only around 20% of human PDAC cases present with actionable genetic alterations making the epigenome and its transcriptional consequences of particular interest.^5^ To elucidate the transcriptional and chromatin accessibility programs associated with chemoresistance and clinical progression at both temporal and single-cell resolution, we conducted a true single-nucleus multiome analysis (RNA and ATAC sequencing) on patient #20. This patient had three organoid lines derived from distinct disease stages and presented as a *KRAS* wild-type case undergoing targeted therapies due to an alternative oncogenic driver (**Figure 5A**). In total 10,408 nuclei from a set of three longitudinal liver metastasis PDO lines (sequentially PDO #20.1, #20.2, and #20.3) were sequenced, followed by multimodal data integration using the weighted nearest neighbor (WNN) approach and dimensionality reduction.^49^ Despite batch integration (Harmony algorithm^50^) of the three distinct samples, cells originating from the same PDO line appeared in closer proximity in the uniform manifold approximation and projection for dimension reduction (UMAP) embedding compared to cells from other samples (**Figures 5B–5D**), suggesting shared biological features or related transcriptional states characteristic for each PDO line. The separation of cells was most pronounced when comparing lines #20.1/2 to #20.3, with the latter being entirely distinct. This observation was consistently supported by transcriptional, open chromatin (OC) level and WNN analyses (**Figures 5B–5D**). Interestingly, PDO #20.1 and #20.2 (derived from sequential biopsies of the same progressive metastatic lesion) positioned closer together in the UMAP embedding, showing partially overlapping OC but distinct transcriptional profiles, underlining the efficacy of our organoid platform in capturing intratumoral molecular heterogeneity. These differences were further reflected in a subsequent GO term enrichment analysis (**Figures 5E–5G**). PDO #20.1 cells were enriched for genes associated with actin organization, cell adhesion, and motility features, in contrast to #20.2 cells, which exhibited decreased association with these biological programs (**Figures 5E and 5F**). This transcriptional shift, concomitant with heightened expression of genes associated with protein and metabolic processes in the PDO line isolated from the second biopsy, may mirror the molecular evolution of cells originating from a shared metastatic source, as well as a direct consequence of the applied therapeutic regimen. The PDO #20.3 transcriptional profile was associated with cell motility, regulation of RNA processing, and neurogenesis – a protumoral program widely described in the context of aggressive PC^51,52^ (**Figure 5G**). To gather further insight into the transcriptional dynamics of longitudinal PDOs and infer single-cell trajectories and cell fate decisions, we conducted a Monocle pseudotime analysis (**Figure 5H**). Our investigation identified two distinct trajectories originating from discrete PDO #20.1-derived cell clusters, transitioning and converging toward a common group of #20.2 cells. This observation suggests the coevolution of distinct transient cell states over time, accompanied by different transcriptional reconfigurations, ultimately leading to the development of a more homogeneous metastatic population. In line with the independent clustering of the PDO #20.3 cells (**Figure 5I**), no dynamic transition from the PDO #20.1 or #20.2 cluster was inferred, thus reinforcing the concept of distinct identities and evolutionary paths for malignant cells derived from spatially separated metastases, as confirmed by clinical documentation (**Figure 5A**). Furthermore, additional unsupervised cell clustering (**Figure 5I**) and gene term overlap analyses (**Figure 5J**), paralleled with the inferred cell trajectories and revealed specific sequential gene expression programing. Cluster 0, serving as the initial compartment for the two inferred cell dynamic trajectories and predominantly comprising PDO #20.1 cells, exhibited enriched gene expression associated with cell motility and adhesion features, consistent with our previous findings (**Figure 5E**). Interestingly, subsequent transition toward less invasive transcriptional programs (cluster 1 to 3), which included transient upregulation of genes associated with amino acid metabolism (cluster 2), were observed in the PDO #20.2 cells (**Figure 5F**). This was followed by the downregulation of cell cycle-related biological programing in cluster 4 (**Figure 5J**). In line with our previous findings, the cell cluster 5, solely composed of PDO #20.3 cells, exhibited heightened gene expression associated with cell motility and transcription regulation, whereas genes involved in NFE2L2 signaling were downregulated (**Figures 5G–5J**). Concomitantly, chromatin accessibility analysis using chromVAR unveiled a marked decline in the accessibility of AP-1 motifs, suggesting a reduced activity of this transcription factor complex across tumor progression stages (**Figures S5A and S5B**). Specifically, we observed a significant downregulation of AP-1, JUN, and JUND target genes in PDO #20.2 compared to PDO #20.1, whereas the PDO #20.3 cells exhibited a decrease in the FOSL1-mediated transcriptomic program relative to PDO #20.2 (**Figure S5C**). Overall, our approach successfully delineated the temporal tumor evolution within a unique patient, thereby emphasizing the value of PDOs as powerful tools for dissecting intricate molecular dynamics inherent in pancreatic cancer progression and metastasis.

**Figure 5.**
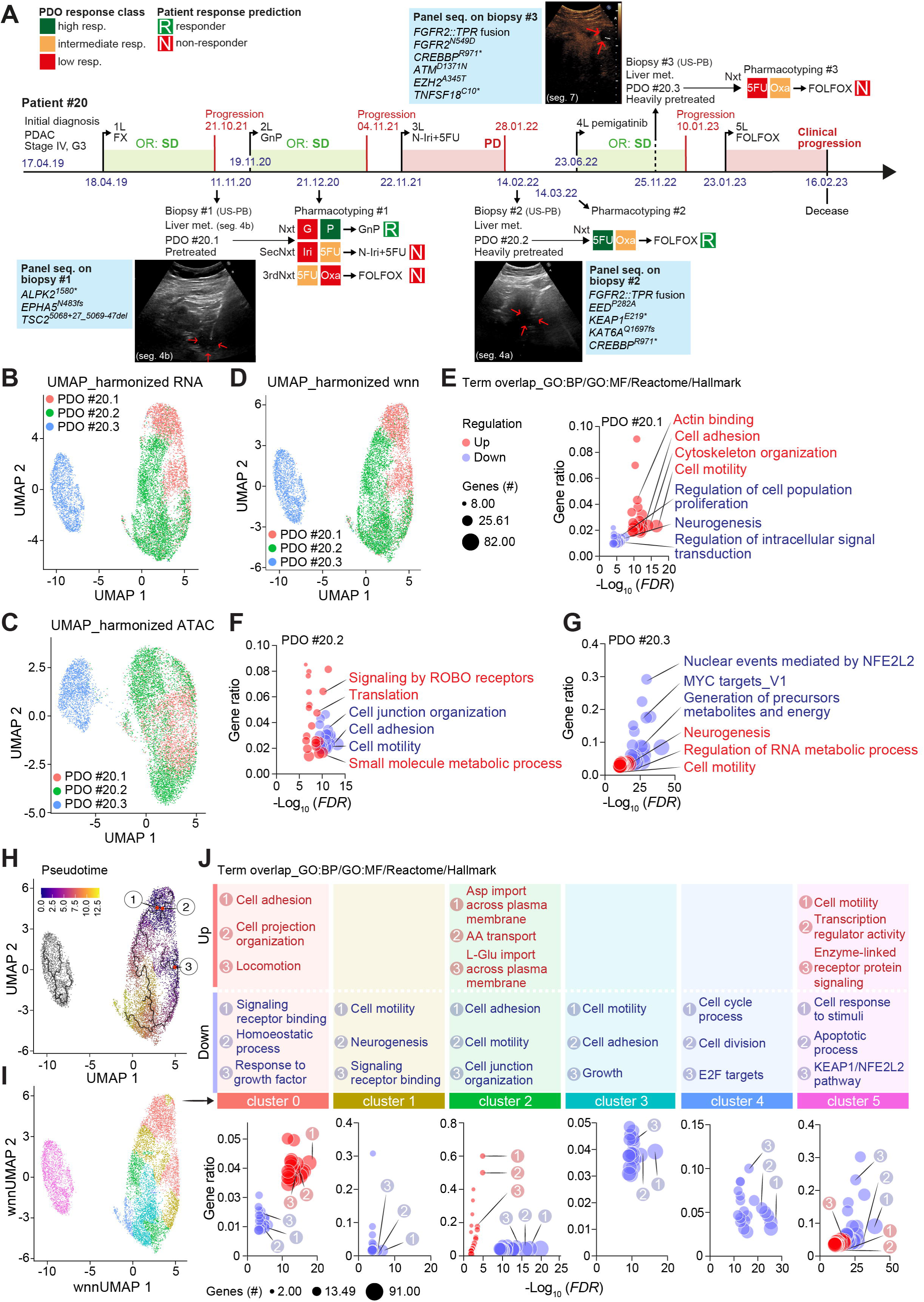
Multiome analysis on a unique longitudinal patient. (A) Patient #20 clinical course. Red arrows point the liver metastases punctured during the ultrasound-guided percutaneous biopsies. Liver metastasis punctured during biopsy #3 is shown under contrast-enhanced ultrasonography. (B–D) UMAP embedding visualizing snRNA (B), snATAC (C), and integrated snRNA and snATAC-seq clusters of cells (D) across the three PDO lines generated from patient #20 as shown in (A). (E–G) Representative GO:BP/GO:MF/Reactome/Hallmark functional annotations for up and downregulated genes in cells from PDO #20.1 (E), PDO #20.2 (F), and PDO #20.3 (G) with fold change ≥2 and p<0.05. (H) UMAP embedding visualizing the inferred position of PDO cells along transcriptomic-based single-cell pseudotime trajectory. (I–J) UMAP embedding visualizing pseudotime-based cell clusters across the three PDO lines (I) and corresponding representative GO:BP/GO:MF/Reactome/Hallmark functional annotations for up and downregulated genes within the six identified cell clusters (J) with fold change ≥2 and p<0.05. 1L, palliative first-line chemotherapy; 2L, palliative second-line chemotherapy; 3L, palliative third-line chemotherapy; 3rdNxt, third next therapy line; 4L, palliative fourth-line chemotherapy; 5FU, 5-fluorouracil; 5L, palliative fifth-line chemotherapy; FOLFOX, folinic acid, 5-fluorouracil, and oxaliplatin; G, gemcitabine; GnP, gemcitabine plus nanoparticle albumin-bound paclitaxel; Iri, irinotecan; n-Iri, nanoliposomal irinotecan; met., metastasis; Nxt, next therapy line; OR, objective response; Oxa, oxaliplatin; P, paclitaxel; PD, progressive disease; PDAC, pancreatic ductal adenocarcinoma; PDO, patient-derived organoid; resp., responder; SD, stable disease; SecNxt, second next therapy line; seg., segment; seq., sequencing; US-PB, ultrasound-guided percutaneous biopsy; wnn, weighted nearest neighbor.

To comprehensively integrate the single-cell sequencing results with the previously acquired genetic and clinical data, we focused on elucidating the FGFR2-related transcriptional program. Indeed, upon disease progression following third-line palliative treatment, a second biopsy was performed to identify putative options of targeted therapies; gene panel sequencing analysis uncovered an activating *FGFR2::TPR* gene fusion, preserving the FGFR2 tyrosine kinase domain (**Figure 6A**). To infer potential FGFR2-related functional molecular network, we first subjected *FGFR2* and the genes significantly enriched in each PDO sample to the STRING database analysis tool and further assessed their expression levels (**Figures 6B–6D and S6A**). Although *FGFR2* expression and chromatin accessibility peaked in organoids derived from the second biopsy (PDO #20.2) (**Figures 6D and 6E**), our analysis inferred distinct FGFR2-related networks in each longitudinal cell cluster (**Figures 6F, 6G, and S6B**). Interestingly, the FGFR2 signaling identified in PDO #20.1 correlated with the upregulation of genes associated with developmental and morphogenic biological processes, while the FGFR2-related networks in PDO #20.2 and PDO #20.3 were more closely linked to protumorigenic signaling pathways. Specifically, genes such as *POLR2L*, *POLR2I*, *PLCB1*, *LIFR*, *STAT3*, and *CCND1*, identified as potentially functional partners to FGFR2 in PDO #20.2, exhibited the highest expression levels compared to PDO #20.1 and PDO #20.3, correlating with FGFR2 signaling in this disease context (**Figures 6D**). Intriguingly, the inferred FGFR2-partner genes, identified via the STRING network in PDO #20.2, were also connected to two gene subclusters involved in protein translation regulation and small molecule metabolism, corroborating our previous findings (**Figures 5F, 5J, and 6C**) and suggesting a central role for FGFR2 signaling in modulating PDO #20.2 transcriptional profile. Finally, PDO #20.3 exhibited the largest number of genes associated with FGFR2 (15 genes: *TIMP1*, *OGA*, *CBLB*, *ERBB3*, *SOX6*, *FN1*, *PTEN*, *NEDD4L*, *PLCG1*, *RPS6KA3*, *GTF2F2*, *GATA6*, *IGF2R*, *FYN*, and *TOX3*). These FGFR2-related genes in PDO #20.3 were further linked to several protumorigenic signaling pathways, including regulation of cell death, proliferation, and signal transduction via growth factor receptors (**Figure 6G**). This particular evolution may be connected to the detection of a FGFR2^N549D^ mutation specifically mediating resistance to FGFR inhibitors,^53^ in biopsy #20.3 collected after fourth-line palliative treatment with pemigatinib (**Figure 5A**). Despite an initial positive clinical response, patient #20 eventually experienced disease progression on this targeted therapy, suggesting that the FGFR2 signaling network identified in #20.3 does reflect specific oncogenic molecular programs driven by the resistance-mediating FGFR2^N549D^ mutation (**Figure 6H**).

**Figure 6.**
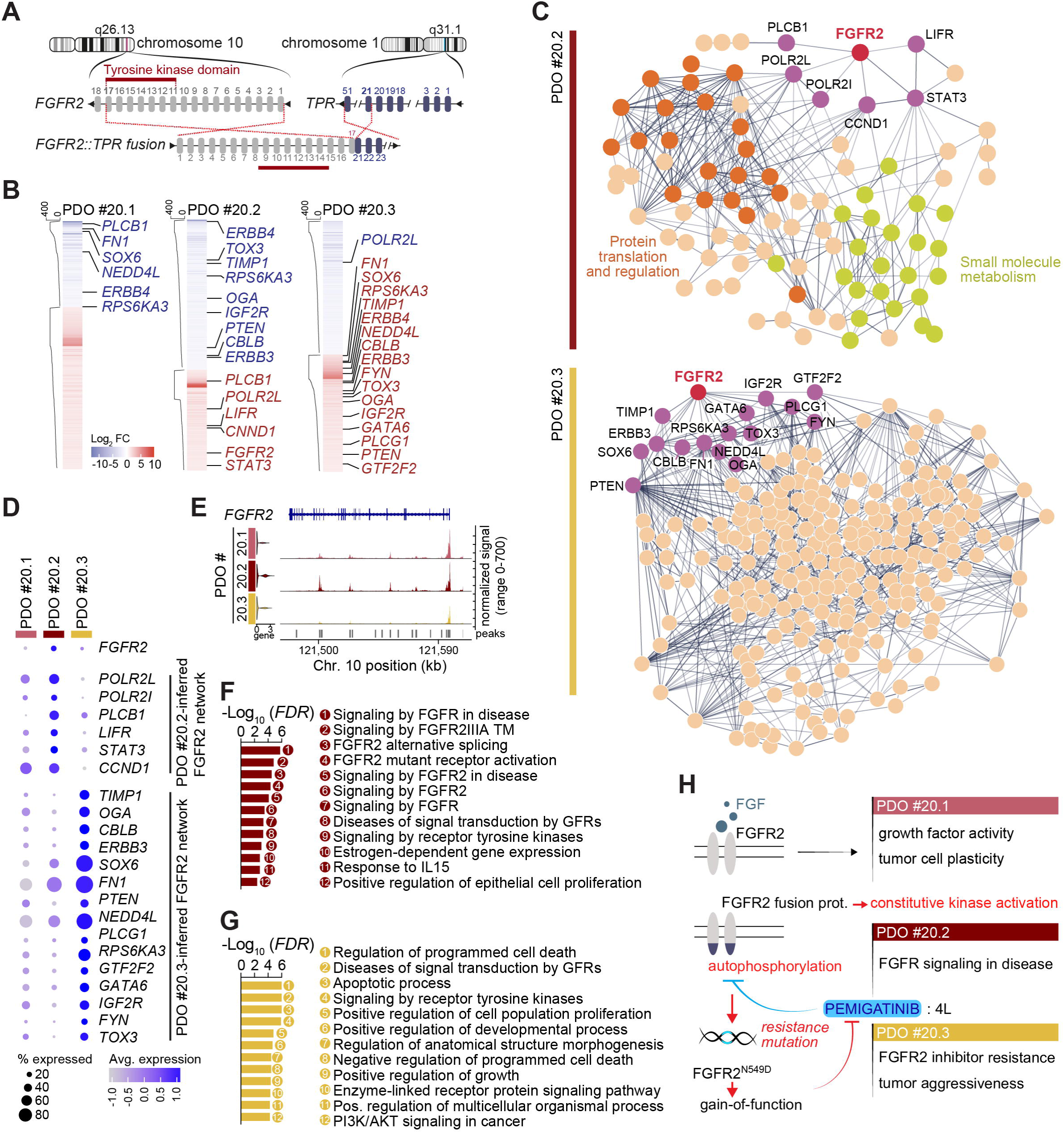
Patient-derived organoids, bona fide systems to investigate therapy-mediated resistance mechanisms. (A) Schematic illustration of the detected *FGFR2::TPR* fusion in patient #20. (B) Heatmap depicting differentially expressed genes in each PDO line with fold change ≥2 and p<0.05. -Log_10_ p-values are plotted. FGFR2-related genes are highlighted. (C) STRING plots depicting inferred molecular FGFR2 network in PDO #20.2 (upper panel) and PDO #20.3 (lower panel) carrying the *FGFR2::TPR* fusion shown in (A). Upregulated genes with fold change ≥2 and p<0.05 were used to generate the STRING plots. (D) Dot plot showing expression of genes inferred as potential FGFR2 partners in (C). Color intensity indicates the average expression for the corresponding gene. Dot size reflects the proportion of cells with non-null expression of the gene. (E) Sequencing tracks showing ATAC-seq peaks of *FGFR2* gene in cells from the three PDO #20 lines. (F–G) Representative GO:BP/GO:MF/Reactome/Hallmark functional annotations in PDO #20.2 (F) PDO #20.3 (G), using genes with fold change ≥2 and p<0.05 depicted in (D). (H) Schematic illustration of the FGF/FGFR2 molecular axis across the disease progression of patient #20, inferred from the single-cell sequencing analysis conducted on corresponding PDO lines. 4L, palliative fourth-line chemotherapy; PDO, patient-derived organoid.

## DISCUSSION

PDOs show significant promise to advance functional precision oncology.^19,54,55^ In recent years, these tumor avatars have been effectively utilized in *in vitro* drug testing to tailor cancer treatments and predict therapy responses. However, the ability to draw overarching conclusions from organoid preclinical studies has been limited by small sample sizes and variations in the scenarios examined.^20,21,24,56^ A recent prospective trial involving a larger cohort of 34 organoid lines alongside corresponding clinical responses demonstrated an impressive 91% efficacy, primarily in the palliative setting.^25^ In the current study, we showcase the potential of PDOs to further personalize therapy within a larger PC cohort comprising 46 organoid lines that matched patient clinical responses. This includes data from 94 lines of therapy across three treatment settings. Importantly, our cohort accurately represents the broader PC population regarding clinical characteristics and mutational profiles.^43,57^

Early therapeutic intervention is vital for PC patients. As observed in our cohort, the median time to treatment initiation (TTT) is usually under a month,^58^ necessitating that PDO pharmacotyping data align with this timeframe – a significant technical challenge. This is especially critical in advanced PC, where first-line therapy must begin urgently, with a median TTT of 14.5 days in our cohort. Unfortunately, a median turnaround time for pharmacotyping of 32.0 days still prevents PDOs from guiding first palliative line treatment in optimal scenarios albeit we optimized assay conditions when compared to other studies.^18,21,24,25^ However, organoid-based prediction following early detection of therapy failures could guide oncologists to promptly switch to a treatment anticipated to be effective. In resectable stages, adjuvant therapy completion is correlated with enhanced patient survival, with generally a TTT above six weeks^59,60^ (59.0 days in our cohort). Additionally, initiation of a neoadjuvant therapy within six weeks post-diagnosis was previously associated with improved survival.^61^ In most cases, PDO profiles were completed in less than six weeks (32 of 40), demonstrating the potential of our organoid model to guide neoadjuvant or adjuvant therapies within a clinically meaningful timeframe. However, the true clinical benefit of PDO-based treatment decisions requires further assessment through prospective randomized clinical trials.^62^ We are currently advancing our efforts with the UNITEPANC trial (AIO-PAK-0424), a prospective interventional study led by the AIO Pancreatic Cancer Group in collaboration with five other medical centers in Germany. This proof-of-concept trial aims to evaluate the performance of organoids in predicting the efficacy of adjuvant treatment for resectable PDACs.

Our study also examined the relationship between pharmacotyping data from a comprehensive PDO registry and clinical outcomes in PC patients. Although this is one of the largest studies conducted, the number of patients available for clinical therapeutic correlation still remains relatively low, compared to the total number of established PDO lines, underscoring the aggressive nature of pancreatic cancer, which often leads to rapid deterioration before treatment initiation or restaging. In the palliative setting, we analyzed responses to 13 therapeutic regimens across more than 80 subsequent chemotherapy lines, finding a strong correlation between organoid sensitivities and clinical responses. Importantly, prior treatment status overall did not significantly affect our match ratio – despite lower accuracy in heavily pretreated organoids – suggesting that PDOs may reliably predict responses across treatment lines, including those following initial therapies. In evaluating (neo)adjuvant outcomes, we observed concordance between organoid and clinical drug responses, indicating that pharmacotyping can inform treatment strategies prior and after surgical interventions. In treatment-naïve patients, the use of predicted effective therapies significantly improved palliative outcomes, demonstrating the clinical relevance of our PDO platform. The overall prediction accuracy of 73.4% attests to its robustness across various treatment scenarios. These findings emphasize the need for innovative functional precision oncology, especially in challenging cases like pancreatic cancer, where conventional methods often fall short.^15^ While our PDO platform showed high accuracy in predicting treatment responsiveness, it also underestimated therapy unresponsiveness, especially with the gemcitabine plus nab-paclitaxel regimen. Current organoid systems have intrinsic limitations due to a lack of stromal components such as cancer-associated fibroblasts, which have been shown to affect drug sensitivity.^63–65^ The absence of these critical components may hinder prediction efficacy, as the tumor–stroma interactions are complex and significant for understanding PC fully.^66,67^ The development of patient-matched multilineage organoids could better represent these interactions and capture resistance mechanisms.^68^ Nevertheless, our approach successfully identified potential treatment vulnerabilities in 96.4% of PDO lines and effective combinations in 87.9%. This starkly contrasts with the 46.4% response rate in patients treated without PDO guidance. Ultimately, our research underscores the significant role that organoid technology could play in improving therapeutic outcomes for patients with this challenging disease.

The promising potential of our PDO model to personalize PC therapy was particularly illustrated by two longitudinal study cases. In patient #18, initially diagnosed with solid pseudopapillary neoplasm and carrying a pathogenic CHEK2^A459T^ mutation, sequential organoid lines were derived from liver lesions before and during olaparib therapy after a late post-resection recurrence. Treatment decision was made as loss-of-function mutations in DNA damage repair pathways may cause vulnerability particularly to DNA-damaging agents as PARP inhibitors.^12,69,70^ Notably, *in vitro* PDO chemoresponses reflected the patient’s prolonged therapeutic response upon olaparib maintenance therapy. The first organoid line demonstrated high susceptibility to olaparib, while the second was classified as an intermediate responder. Interestingly, the clonal evolution suggested drug-induced selection of a pre-existing olaparib-resistant subclone in PDO #18.2. Despite this indication of a potentially altered sensitivity, the patient remained stable during olaparib treatment indicating overall disease control.

The disease course of patient #20, who presented a unique clinical trajectory with KRAS wild-type status, illustrates the limitations of relying solely on genomic sequencing in PC, and underscores the need to systematically explore the epigenome and transcriptome for comprehensive understanding of tumor biology and chemoresistance. Using DNA, RNA, and ATAC-sequencing across multiple disease stages represented by distinct PDO lines, we captured the complex genetic, transcriptional, and chromatin accessibility programs associated with treatment response and resistance. Notably, we identified an *FGFR2::TPR* fusion transcript which allowed us to examine FGFR2-related transcriptional programs throughout the treatment process. After three lines of standard chemotherapy, the identification of this actionable fusion enabled successful targeted therapy with pemigatinib, resulting in an extended response. However, a third biopsy later revealed a drug-driven resistance-mediating *FGFR2* mutation after five months of treatment, alongside new pathogenic genomic alterations, underpinning the value of genomic information to functional treatment response.

Our comprehensive multiome analysis revealed specific cellular responses and oncogenic molecular programs at each stage, illustrating a clear evolutionary trajectory from PDO #20.1 to PDO #20.2, with no connection to PDO #20.3. The distinct gene networks identified within the PDO lines underscore the adaptive nature of PC under therapeutic pressure. Notably, PDO #20.3 displayed heightened expression of genes associated with tumorigenesis and resistance, even as initial treatments proved effective. This finding is consistent with existing literature on the connection between genetic alterations and treatment responses in PC, emphasizing the necessity of integrating genomic, transcriptomic, and epigenomic profiling.^19,71^ Furthermore, the single-cell resolution captured the emergence of rapidly-evolving clones in response to tailored therapy pressures.

In conclusion, our study underscores the significant potential of PDOs in advancing precision oncology for PC. Our unique living organoid biobank encompasses a wide range of PC stages and therapeutic scenarios, all managed within an organoid core facility that centralizes specialist expertise and ensures rigorous data quality monitoring and adhesion to best research practices. This was illuminated by the elaboration of a highly reliable robot-assisted pharmacotyping procedure. In the future, engineering efforts will enable the development of complex organoid avatars that more accurately replicate individual tumor characteristics. Importantly, enhancing the reconstruction of cell-extrinsic chemoresistance interactions will further refine this system. Identifying actionable events in PC is crucial, requiring sometimes comprehensive analyses of complex genetic, transcriptional, and chromatin accessibility data to capture the diverse cellular processes underlying therapy response and resistance. Moving forward, we advocate for further validation through prospective randomized clinical trials, which will be pivotal in establishing PDOs as standard practice in treatment decisions and improving therapeutic outcomes for patients facing this challenging disease.

## ACKNOWLEDGMENTS

Main funding was provided by the Hans Beger Foundation and the Deutsche Krebshilfe (German Cancer Aid) grants 70114761 to A.K. and 70115292 to L.P.. A.K. is also funded by the “Heisenberg-Programm” KL 2544/6-1. Additional funding to L.P. came from Deutsche Forschungsgemeinschaft (DFG) PE 3337/1-1. E.R. received funding by the Bausteinprogramm of Ulm University (L.SBN.0193). E.R. is a Hertha-Nathorff-Programm fellow. A.K. and T.S. are speakers of an Else Kröner Research School for Physicians. We are deeply grateful to Kuhn Elektro-Technik GmbH for supporting our research to fight pancreatic cancer. We would like to thank Katrin Jochmann, Katrin Köhn, and Natalie Paul for their essential experimental support. The authors especially thank Tapan Joshi and Verena Renz for their outstanding technical support. The authors thank Dr. Ninel Azoitei for helpful scientific exchange. We would like to thank the Core Facility Organoids of the Medical Faculty at Ulm University for providing support and instrumentation.

## AUTHOR CONTRIBUTIONS

**J.G.:** Conceptualization, methodology, formal analysis, investigation, writing–original draft, visualization, funding acquisition. **L.P.:** Conceptualization, methodology, formal analysis, resources, writing–original draft. **J.L.:** Conceptualization, methodology, formal analysis, investigation, resources. **Y.J.R.:** Conceptualization, methodology, formal analysis, resources. **J.D.S.:** Methodology, formal analysis, investigation. **J.M.K.:** Methodology, formal analysis, investigation. **E.R.:** Formal analysis, writing–original draft, visualization. **D.S.:** Methodology, formal analysis, investigation. **E.Z.:** Formal analysis, investigation. **B.M.:** Formal analysis. **R.M.:** Resources. **N.T.G.:** Resources. **J.K.K.:** Resources. **M.H.:** Resources. **T.J.E.:** Resources. **K.M.D.:** Resources. **A.K.B.:** Formal analysis, Resources. **B.K.:** Resources. **C.M.:** Resources. **A.L.M.:** Resources. **N.N.R.:** Resources. **M.K.:** Resources. **H.A.K.:** Methodology, resources, supervision. **T.S.:** Resources, patient recruitment, project administration. **A.K.:** Conceptualization, investigation, resources, writing–original draft, supervision, project administration, funding acquisition.

## STAR METHODS

### Patient selection and ethics statement

Patients with a presumed or confirmed diagnosis of PC, regardless of tumor stage or treatment status, were eligible to participate in the research project at the University Hospital Ulm. Patients with PC suspicion underwent routine diagnostic work-up and tissue acquisition to confirm the diagnosis or surgical resection independently of their participation to the study. Pretreated patients who were re-biopsied for participation in a clinical study or for panel-sequencing, were also included in this study. Exclusion criteria were inability to provide written informed consent or lack of clinical indication for invasive tissue acquisition. Patient enrollment in the study was conducted between July 2019 and April 2024 at the University Hospital Ulm. The project was approved by the local ethics committee (project number 72/19), and written informed consent was obtained from all patients.

### Tumor specimen collection

All ultrasound-guided procedures were performed at the Department of Internal Medicine 1 of the University Hospital Ulm. One or two tumor specimens (sampled with a 16 or 18-gauge needle for ultrasound-guided biopsies of the primary tumor and liver metastases; a standard 19-gauge needle for endoscopic ultrasound-guided biopsies) were processed for organoid isolation. Surgical specimens were received intraoperatively from the Department of General and Visceral Surgery of the University Hospital Ulm. Samples were placed into advanced DMEM medium supplemented with 100 μg/mL primocin (InvivoGen) and 10 mg/mL bovine serum albumin (Sigma-Aldrich), and were kept on ice until isolation (within 30 min to 3 h). Diagnosis of pancreatic cancer was confirmed by routine pathology assessment.

### Clinical evaluation of treatment response

Patients were treated based on their previous treatment history, actionable mutations (if available), and performance status, independently of our study. Treatment response was routinely evaluated by computed tomography scan per clinical practice standard. In the palliative setting, clinical evaluation of treatment response was assessed according to RECIST v1.1 (response evaluation criteria in solid tumors).^31^

### Isolation and culture of patient-derived organoids

Briefly, tissue biopsy samples and surgical tumor pieces were minced into 1 mm fragments with a sterile scalpel and enzymatically digested with accutase (Sigma-Aldrich) for 40 min at 37 °C. Cells were then seeded into Matrigel Growth Factor Reduced (GFR) (Corning) domes covered with organoid culture medium containing WNT3A and RSPOI-conditioned media.^72^ Surgical specimens were digested with collagenase II (Sigma-Aldrich) and strained before seeding in Matrigel GFR. Organoids were propagated at 37 °C under 5% (v/v) CO_2_ atmosphere and passaged for cell line expansion every 5 to 14 days depending on cell density and growth rate. Mycoplasma tests were regularly performed using the MycoSPX PCR kit (Biontex). WNT signaling pathway activity of the WNT3A-conditioned medium was assessed with a dual luciferase (Firefly-Renilla, BPS Bioscience) assay system (TCF/LEF reporter kit, BPS Bioscience). All experiments were performed between passage 4 and 25.

### Organoids imaging

Brightfield images were acquired using a 2× or 4× objective mounted on a Keyence BZ-X810 microscope (Keyence) using the BZ-X800 Viewer software (Keyence).

### Pharmacotyping

The miniaturized robot-assisted pharmacotyping procedure is described in **Document S1**. Chemotherapeutics were tested in triplicates in ten concentrations covering four orders of magnitudes (13 nM to 50 μM). Percentage of cell viability was determined using this formula: (S2-S1)_Drug_/(S2-S1)_Ctrl_. The area under the curve (AUC) was estimated using the trapezoidal rule with Prism v8 (GraphPad Software). For each single chemotherapeutic substance, AUCs from the PDO library were classified into three subgroups (low, intermediate, and high responder) using the Jenks natural breaks classification method.^18,24^

5-fluorouracil (NSC 19893), cisplatin (NSC 119875), erlotinib (CP-358774), gemcitabine (LY188011), irinotecan (CPT-11), olaparib (AZD2281), oxaliplatin (L-OHP), paclitaxel (NSC 125973), and palbociclib (PD-0332991) were purchased from Selleckchem.

### Library preparation and next generation sequencing

For library preparation 80 ng of isolated DNA was used in conjunction with the Tumor Mutational Burden panel (# DHS-6600Z, Qiagen) using the QIASeq protocol according to the instructions of the manufacturer. The resulting library was quantified by Qubit measurement using the Qubit dsDNA High Sensitivity Assay kit (Thermo Fisher Scientific) and quality was checked by tape station (HS D1000 ScreenTape assay, Agilent). Subsequently, the library was sequenced on a NovaSeq device (Illumina) and the resulting fastq files were processed using the CLC Genomics Workbench version 23.0.3 (Qiagen). Variants were filtered using following parameter: reference allele: no, EUR 1000GENOMES-phase_3_ensembl_v106_hg38_no_alt_analysis_set: ≤1%, variant allele frequency: ≥3%. All remaining variants were manually curated using the BAM files in conjunction with the integrative genomics viewer (v2.8.13) and classified using the dbSNP, ClinVar, OncoKB, CKB, and Civic databases.

### Whole exome sequencing

#### Sequencing alignment

For read mapping and variant detection, the nf-core^73^ pipeline sarek^74,75^ was applied with the default parameters (Nextflow v23.04.2, nf-core/sarek v3.3.2). FastQC (v0.11.9) and fastp (v0.23.4) were used for quality control and trimming of reads. Reads were mapped to the human reference genome GRCh38 using Burrows-Wheeler Aligner BWA-MEM (v0.7.17-r1188). With Genome Analysis Toolkit GATK (v4.4.0.0), duplicates were tagged, and base quality scores were recalibrated. Samtools (v1.17) and tabix (v1.12) were used for file format conversion and indexing.

#### Detection of somatic variants and mutational burden

Variant Calling was performed on the GRCh38 genome with GATK (v4.4.0.0) in the MuTect2 mode, followed by variant annotation using SnpEff (v5.1d). Both steps were applied within the sarek framework. Tumor mutational burden was calculated by applying pyTMB, a Python tool provided on GitHub by the Institut Curie (https://github.com/bioinfo-pf-curie/TMB).

#### Clonal tracking

To identify copy number alterations and clonal tracking, SuperFreq (v1.4.5)^48^ was applied with default parameters (https://github.com/ChristofferFlensburg/superFreq). Germline samples were used as reference normal samples as required by the pipeline.

### Multiome sequencing

#### Preprocessing

As preprocessing for the single-cell multiome data, poor quality cells were excluded, such as cells with <1,000 or >25,000 RNA molecules detected (as they are potentially empty droplets or duplets). Cells were also discarded if the mitochondrial gene percentage was above 20. Regarding ATAC sequencing, cells with <500 and >70,000 molecules were excluded (again to remove empty droplets and duplets). After preprocessing, the three datasets (PDO #20.1, PDO #20.2, and PDO #20.3) were merged individually for RNA and ATAC sequencing assays. RNA-sequencing data was natural log transformed and normalized for scaling the sequencing depth to 1,000 molecules per cell (Seurat R-package, v5.0.0). ATAC-sequencing data was normalized via term frequency inverse document frequency (TF-IDF) normalization (Seurat R-package, v5.0.0^76^). To remove batch effects between the different samples, the Harmony algorithm was applied (Harmony R-package, v1.1.0^50^) on the data from both modalities individually.

#### Integration of scRNA and scATAC data

Data integration was done with the Seurat-R-package (v5.0.0) using the weighted-nearest neighbor approach (WNN, function FindMultiModalNeighbours^49^) based on the harmony-corrected principle components of scRNA (dimensions/principle components 1 to 11) and scATAC data (dimensions/principle components 2 to 50). For visualization, 2D-UMAP plots were created for harmonized scRNA, scATAC, and WNN-integrated multiome-data.

#### Finding markers for clusters

Markers for the individual cells from PDO #20.1, PDO #20.2, and PDO #20.3 were identified using differential gene expression analysis. Therefore, differential expression analysis was performed on the cells of each cluster against the remaining cells (function FindAllMarkers in Seurat, v5.0.0).

#### Pseudotime trajectories

For the computation of pseudotime trajectories, the WNN-integrated data was transferred from Seurat (v5.0.0) to monocle3 (v1.3.4) using the R-package Seurat Wrappers (v0.3.2). Cells were sub-clustered using the Louvain clustering approach with a resolution of 0.001 (function cluster_cells in monocle3). Pseudotime trajectories were computed using the resulting sub-clustering (function learn_graph in monocle3^77^). To determine marker genes along the pseudotime trajectory, the pseudotime values for each cell were binned in four intervals (0-3,4-6,7-9,10-12 + Inf for the disconnected cluster). Cells were grouped according to these intervals, and differential gene expression analysis was performed for all cells in each interval versus all other cells (findAllMarkers function in Seurat).

#### Peak analysis in scATAC data

Peak analysis was done using R-Package Signac (v1.12.9000^78^). GC content, region lengths, and dinucleotide base frequencies for regions were computed (RegionStats function) using GRCh38 as reference genome (R-package BSgenome.Hsapiens.UCSC.hg38). ATAC peaks, which are correlated with the expression of nearby genes, were searched (LinkPeaks function). Coverage plots show the frequency of Tn5 insertion events for the different clusters PDO #20.1, PDO #20.2, and PDO #20.3 individually. Enrichment scores of motifs in accessible regions were calculated using chromVAR.^79^

#### Gene term enrichment analyses

Gene term enrichment analyses were conducted using the Molecular Signatures Database (MSigDB) with the Gene Ontology, Reactome, and Hallmark datasets. Significant enrichments were defined with *FDR*<0.05. Interaction networks were generated using the Search Tool for the Retrieval of Interacting Genes/Proteins (STRING) v12.0 (interaction score set to medium confidence).

### Data and code availability

Raw genomic sequencing data generated in this study are not publicly available to ensure protection of patient personal data and privacy. Raw snRNA-seq and ATAC-seq data generated for this project is available on GEO with accession number “-“ (reviewer secure token:-).

### Statistical analysis

Prism (GraphPad) software was used for statistical analysis and graphical representation of the data. Correlation analyses were conducted using Pearson’s correlation test. Statistical significances in comparing frequency distributions were tested using Fisher’s exact test. For overall and progression-free survival comparisons, statistical significances were tested using log-rank (Mantel-Cox) test. All tests were considered to be statistically significant when p<0.05.

## SUPPLEMENTARY FIGURE LEGENDS

**Figure S1. Related to Figure 1. Organoid line derivation efficacies**

(A) Kaplan-Meier analysis showing overall survival of patients with stage I(A+B), II(A+B), III, and IV pancreatic cancer (PC).

(B–C) Pie charts depicting patient-derived organoid line derivation efficacies according to biological material origin (B) and treatment status (C).

(D) Kaplan-Meier analysis showing overall survival of pancreatic cancer (PC) patients with successful or failed line derivation.

(E) Comparison of time between biopsy and decease or lost to follow-up for PC patients with successful or failed line derivation.

**Figure S2. Related to Figure 2 and 3. Patient-derived organoids, preclinical avatars to predict patient therapeutic response in pancreatic cancer**

(A–B) Time to organoid pharmacotyping according to used sampling method (A) and tumor grade and stage (B).

(C) Time to treatment initiation, first restaging CT scan, and progression upon palliative first-line therapy.

(D) Pearson correlation between individual organoid response to all screened single agents.

(E) Swimmer plot showing progression-free survival of PC patients upon palliation who received consecutive regimens predicted to be effective or ineffective by our PDO pharmacotyping-based model, and corresponding progression-free survival ratios.

(F) Progression-free survival ratios of the cohort of PC patients upon palliation who qualified for ≥2 therapy lines.

(G) Hit identification rate for a SOC agent.

(H) Therapeutic lines analyzed in the study.

1L, palliative first-line chemotherapy; 2L, palliative second-line chemotherapy; 3L, palliative third-line chemotherapy; 4L, palliative fourth-line chemotherapy; 5FU, 5-fluorouracil; 5L, palliative fifth-line chemotherapy; 6L, palliative sixth-line chemotherapy; Cape, capecitabine; Cis, cisplatin; Doce, docetaxel; Erlo, erlotinib; EUS-FNB, endoscopic ultrasound-guided fine-needle biopsy; FOLFIRI, folinic acid, 5-fluorouracil, and irinotecan; FOLFIRINOX (folinic acid, 5-fluorouracil, irinotecan, and oxaliplatin); FOLFOX, folinic acid, 5-fluorouracil, and oxaliplatin; G, gemcitabine; GnP, gemcitabine plus nanoparticle albumin-bound paclitaxel; n-Iri, nanoliposomal irinotecan; neg.c., negative correlation; OFF, oxaliplatin, 5-fluorouracil, and folinic acid; Ola, olaparib; Oxa, oxaliplatin; P, paclitaxel; PFS, progression-free survival; Pr, pretreated; SOC, standard-of-care; US-PB, ultrasound-guided percutaneous biopsy.

**Figure S3. Related to Figure 4. Clinical courses of longitudinal patients**

(A–F) Clinical courses of longitudinal patients #4 (A), #10 (B), #13 (C), #39 (D), #40 (E), and #18 (F).

1L, palliative first-line chemotherapy; 2L, palliative second-line chemotherapy; 3L, palliative third-line chemotherapy; 3rdNxt, third next therapy line; 4L, palliative fourth-line chemotherapy; 5FU, 5-fluorouracil; Cape, capecitabine; Curr, current therapy line; Doce, docetaxel; EUS-FNB, endoscopic ultrasound-guided fine-needle biopsy; FOLFOX, folinic acid, 5-fluorouracil, and oxaliplatin; FX, FOLFIRINOX (folinic acid, 5-fluorouracil, irinotecan, and oxaliplatin); G, gemcitabine; GnP, gemcitabine plus nanoparticle albumin-bound paclitaxel; Iri, irinotecan; n-Iri, nanoliposomal irinotecan; met., metastasis; Nxt, next therapy line; Ola, olaparib; OR, objective response; Oxa, oxaliplatin; P, paclitaxel; PD, progressive disease; PDAC, pancreatic ductal adenocarcinoma; PDO, patient-derived organoid; PR, partial response; SD, stable disease; SecNxt, second next therapy line; seg., segment; seq., sequencing; SPN, solid pseudopapillary neoplasms of the pancreas; unspec., unspecified; US-PB, ultrasound-guided percutaneous biopsy.

**Figure S4. Related to Figure 4. Somatic alterations identified within the patient cohort**

(A) Overview of the somatic mutations found in enrolled pancreatic cancer (PC) patients by gene panel sequencing and whole exome sequencing.

(B) Proportions of non-longitudinal PC patients whose tumors harbor *KRAS*, *TP53*, and actionable mutations.

Met., metastasis; N/A, not available; PDO, patient-derived organoid; PT, primary tumor; St, stable; TMB, tumor mutation burden.

**Figure S5. Related to Figure 5. Multiome analysis on a unique longitudinal patient**

(A) chromVAR motif enrichment (enrich.) scores for identified transcription factors in the three PDO #20 lines.

(B) UMAP embedding visualizing *FOS*, *JUN*, *ATF2*, *FOSL1*, *FOSL2*, and *FOS::JUN* TF motif enrichment in cells from the three PDO #20 lines.

(C) AP1 gene sets-based GSEA conducted on transcriptomic data from PDO #20 lines. PDO, patient-derived organoid; wnn, weighted nearest neighbor.

**Figure S6. Related to Figure 6. Patient-derived organoids, bona fide systems to investigate therapy-mediated resistance mechanisms**

(A) STRING plot depicting inferred molecular FGFR2 network in PDO #20.1. Upregulated genes with fold change ≥2 and p<0.05 were used to generate the STRING plots.

(B) Representative GO:BP/GO:MF/Reactome/Hallmark functional annotations in PDO #20.1, using genes with fold change ≥2 and p<0.05.

PDO, patient-derived organoid.

