## Supplementary material for "Pancreatic cancer patient-derived organoids capture therapy response and tumor evolution": Document_S1

**Document S1. Procedure for miniaturized robot-assisted pharmacotyping of pancreatic cancer patient-derived organoids.**

*Seeding cells for pharmacotyping*

Prior to the conduction of this procedure, the pancreatic cancer patient-derived organoids have to be propagated to reach the required biomass of 210,000 cells (500 cells per well) for a 9-drug screening (ten doses in triplicates).

1. Prewarm black 384-well plates labeled with ID stickers at 37 °C, till step 26 (cell seeding for the assay).
2. Carefully remove the culture medium above the cells, from the wall of the well, using the vacuum pump with glass sterile Pasteur pipettes. Do not perturb the GFR-Matrigel (Corning) domes.
3. Apply 500 µL of passaging medium (see table below) to each organoid-containing well.
4. Incubate the cell culture plate for 2.5 hours in a 37 °C incubator under 5% (v/v) $CO_2$ atmosphere. Check under a microscope if the GFR-Matrigel dome is dissolved and if the cells are detached. *NB: the GFR-Matrigel dome should be easily dissolve by pipetting up and down with a 1,000 µL micropipette. If not, the incubation time can be extended to 3 hours.*
5. Add 500 µL of tempered (RT) resuspension medium (see table below) to each well.
6. Resuspend the organoids and GFR-Matrigel by pipetting up and down five times with a 1,000 µL micropipette.
7. Transfer cell suspensions into a 15 mL tube.
8. Flush each well with 500 µL of resuspension medium (see table below) to wash out the remaining cells.
9. Transfer the cell suspension to the corresponding 15 mL tube.
10. Centrifuge at 500 *g* for 5 minutes at RT.
11. Carefully discard the supernatant using the vacuum pump with glass sterile Pasteur pipettes.
12. Carefully resuspend the pellet in 2 mL of PBS with a 1,000 µL micropipette.
13. Centrifuge at 500 *g* for 5 minutes at RT.
14. Carefully discard the supernatant using the vacuum pump with glass sterile Pasteur pipettes.
15. Resuspend the pellet in 500 µL of accutase solution (Sigma-Aldrich) supplemented with 10 µM Rho kinase inhibitor Y-27632, using a 1,000 µL micropipette. *NB: if the cell pellet is huge, the volume of* accutase *solution can be increased up to 1,000 µL.*
16. Gently pipette up and down ten times using a 1,000 µL micropipette.
17. Incubate the cell suspension for 30 minutes in a 37 °C water bath. Every 10 minutes, pipet the suspension up and down ten times using a 1,000 µL micropipette.
    *NB1: this step is critical to obtain a single cell suspension, which is absolutely crucial for the drug response test assay.*
    *NB2: check whether single cells have been obtained via observing the cell suspension in the tube under a microscope. If not yet, extend the accutase incubation up to 60 minutes.*
18. Add 500 µL of tempered (RT) resuspension medium to inhibit the accutase.
19. Pipette the cell suspension up and down ten times using a 1,000 µL micropipette.
20. Retrieve 20 µL of cell suspension and dilute it 1:1 in trypan blue.
21. Count living cells in a hematocytometer (trypan blue exclusion method).
22. Transfer two times 105,000 cells in a 1.5 mL tube.
23. Centrifuge at 500 *g* for 5 minutes at RT. *NB: a small cell pellet is observable.*
24. Carefully discard the supernatant using the vacuum pump with glass sterile Pasteur pipettes, then with a 200 µL micropipette, without disrupting the pellet.

25. Resuspend each pellet in 210 µL of GFR-Matrigel, by pipetting up and down ten times using a 1,000 µL micropipette. *NB: avoid drawing air bubbles into the suspension.*
26. Place the prewarmed 384-well plate in the Mantis liquid dispenser and insert the tip with the cells into a high-volume chip. Proceed with the cell seeding using the Mantis liquid dispenser (Formulatrix). *NB: to avoid edge effect due to medium evaporation, do not seed cells in edge wells (rows A, B, O, and P and columns 1, 2, 23, and 24; this will leave 240 usable wells per plate).*
27. Check the plate for empty wells. Make sure that domes are properly at the bottom of the wells and not to the sides. *NB: document wrongly processed wells.*
28. Incubate the 384-well plate for 15 minutes in a 37 °C incubator under 5% (v/v) $CO_2$ atmosphere.
29. Prepare the required volume (25 µL per well) of human feeding medium (see table below) supplemented with 10 µM Rho kinase inhibitor Y-27632.
30. Apply 25 µL of Y-27632-supplemented human feeding medium into the wells using the Mantis liquid dispenser and the continuous flow chip.
31. Fill edge wells with 50 µL of PBS using the Mantis liquid dispenser.
32. Propagate cells for one day at 37 °C under 5% (v/v) $CO_2$ atmosphere.

*Drug testing*

A broad testing range of ten concentrations, covering four orders of magnitude from micromolar to nanomolar (50.000 µM, 20.000 µM, 8.000 µM, 3.200 µM, 1.280 µM, 0.512 µM, 0.205 µM, 0.082 µM, 0.033 µM, and 0.013 µM) is used in the procedure. All experimental points are performed in triplicates.

33. Thaw the drugs to be tested. *NB: oxaliplatin and cisplatin solutions have to be prepared freshly.*
34. Prepare two drug dilutions of 250.0 µM and 2.5 µM in Rho kinase inhibitor Y-27632-free human-complete-feeding-medium. Include vehicle concentration controls (highest dose of drug solvents). *NB: vortex thoroughly drug stock solutions and working mixes before use.*

Final drug dilution will be performed in the wells using the Mantis liquid dispenser (according to the dilution table below).

| Drug dilution # | Final working concentration (µM) | Volume of CPLT (µL) | Volume of 250.0 µM drug solution (µL) | Volume of 2.5 µM drug solution (µL) |
|---|---|---|---|---|
| 1 | 50.000 | 15.0 | 10.0 | - |
| 2 | 20.000 | 21.0 | 4.0 | - |
| 3 | 8.000 | 23.4 | 1.6 | - |
| 4 | 3.200 | 24.4 | 0.6 | - |
| 5 | 1.280 | 24.7 | 0.3 | - |
| 6 | 0.512 | 14.8 | - | 10.2 |
| 7 | 0.205 | 20.9 | - | 4.1 |
| 8 | 0.082 | 23.4 | - | 1.6 |
| 9 | 0.033 | 24.3 | - | 0.7 |
| 10 | 0.013 | 24.7 | - | 0.3 |
| Vehicle | - | 15.0 | 10.0 (vehicle stock solution) | |

35. Apply the required volumes of medium per well.
36. Add the vehicle controls on cells.
37. Add the drug solutions on cells. Continuous-flow for medium, HV chip for DMSO, LV chip for drug dilution 250.0 µM, LV chip for drug dilution 2.5 µM.
38. Propagate cells for the next four days at 37 °C under 5% (v/v) $CO_2$ atmosphere.

*Measuring cytotoxicity*

Drug cytotoxicity is evaluated by measuring relative dead cell number, with the dead-cell protease assay CytoTox-Glo (Promega), following the manufacturer's protocol and using the Mantis liquid dispenser. Luminescence is measured using a Tecan Infinite M200 Pro (Tecan).

*Required reagents, solutions, and media*

**Passaging medium**

| Reagent | Final concentration | Supplier |
|---|---|---|
| Advanced DMEM/F-12 | - | Life Technologies |
| Collagenase/dispase | 1 mg/mL | Roche |
| Rho kinase inhibitor (Y-27632) | 10 µM | StemCell Technologies |
| Primocin | 100 µg/mL | Invivogen |

Filter the solution using a 0.22 µm filter.

**Human BSA wash medium (Baker et al., 2019)**

| Reagent | Final concentration | Supplier |
|---|---|---|
| DMEM | - | Life Technologies |
| Primocin | 100 µg/mL | Invivogen |
| Bovine serum albumin | 10 mg/mL | Sigma-Aldrich |

Filter the solution using a 0.22 µm filter.

**Human wash medium (Baker et al., 2019)**

| Reagent | Final concentration | Supplier |
|---|---|---|
| Advanced DMEM/F-12 | - | Life Technologies |
| HEPES | 10 mM | Lonza |
| GlutaMAX | 1:100 | Thermo Fisher Scientific |
| Primocin | 100 µg/mL | Invivogen |

Filter the solution using a 0.22 µm filter.

**Human feeding medium (Baker et al., 2019)**

| Reagent | Final concentration | Supplier |
|---|---|---|
| Human wash medium | - | See above |
| WNT3A CM | 1:2 | Conditioned medium |
| R-SPOI CM | 1:10 | Conditioned medium |
| B27 supplement | 1:50 | Thermo Fisher Scientific |
| Nicotinamide | 10.00 mM | Sigma-Aldrich |
| N-acetylcysteine | 1.25 mM | Sigma-Aldrich |
| hEGF | 50 ng/mL | Peprotech |
| hFGF10 | 100 ng/mL | Miltenyi Biotec |
| hGastrin I | 10 nM | Tocris |
| mNoggin | 100 ng/mL | Peprotech |
| A 83-01 | 500 nM | Tocris |
| Primocin | 100 µg/mL | Invivogen |

Filter the solution using a 0.22 µm filter.
