## Supplementary figures and images for "Pancreatic cancer patient-derived organoids capture therapy response and tumor evolution"

### Supplemental Figure S1

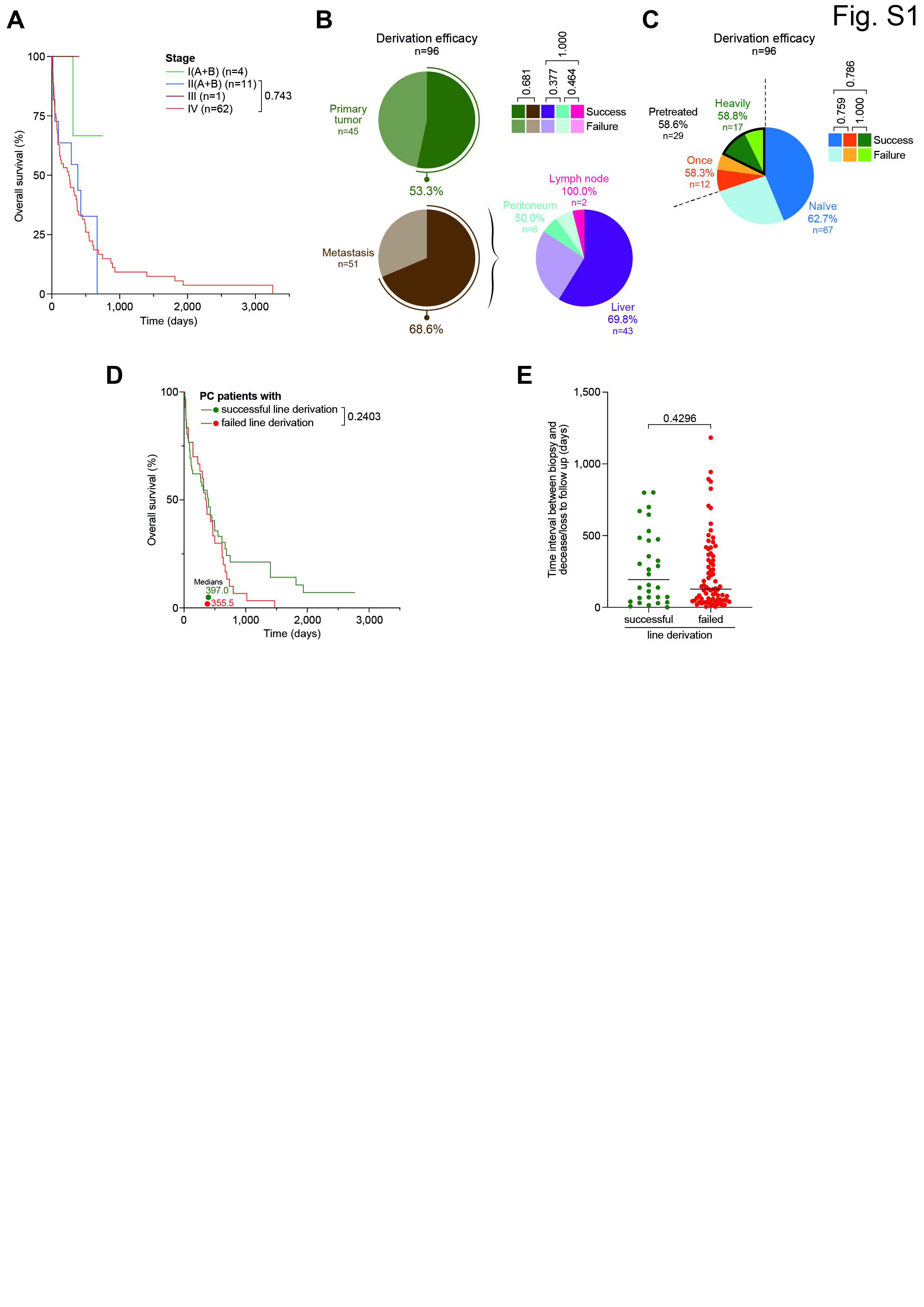

### Supplemental Figure S2

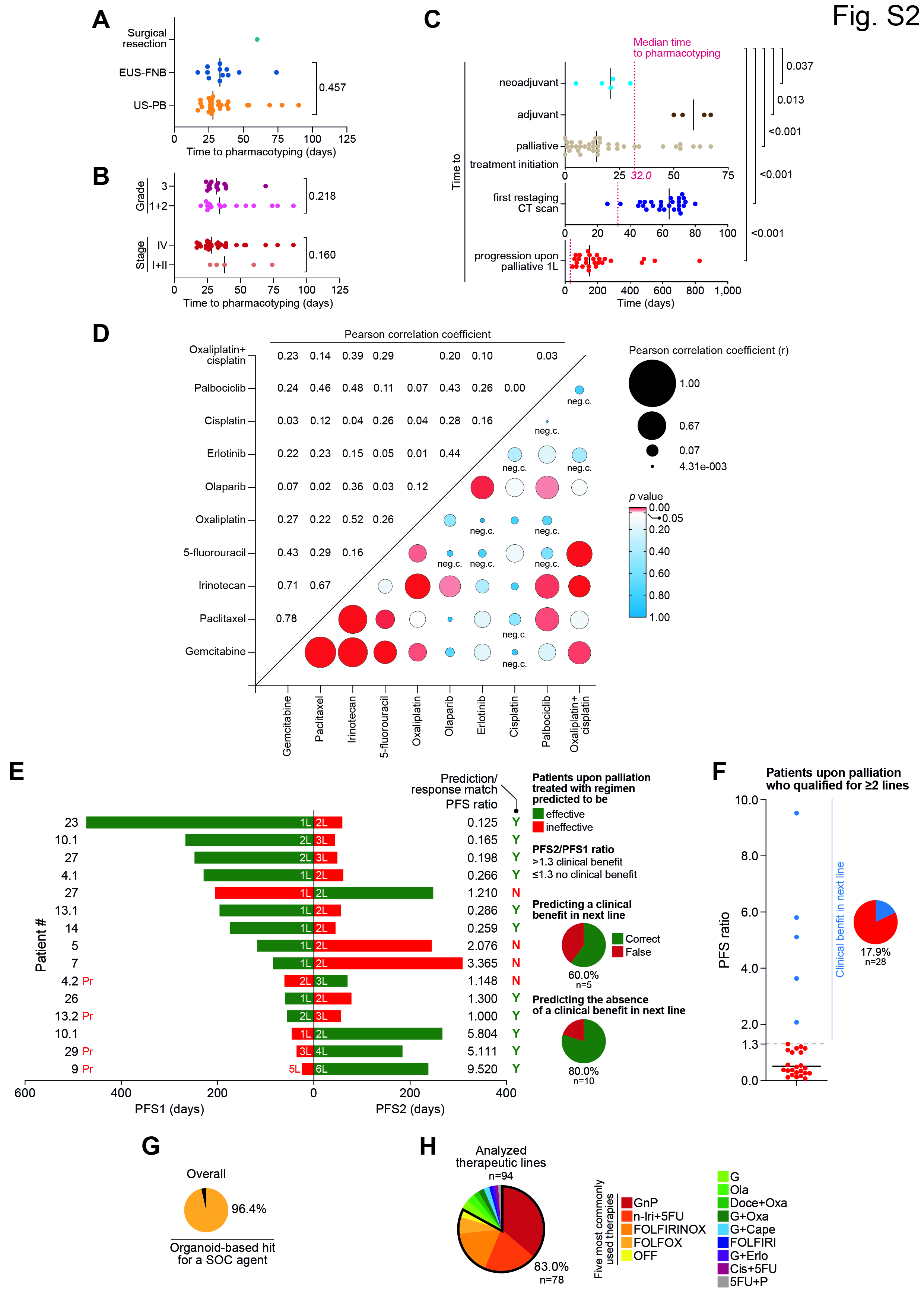

### Supplemental Figure S3

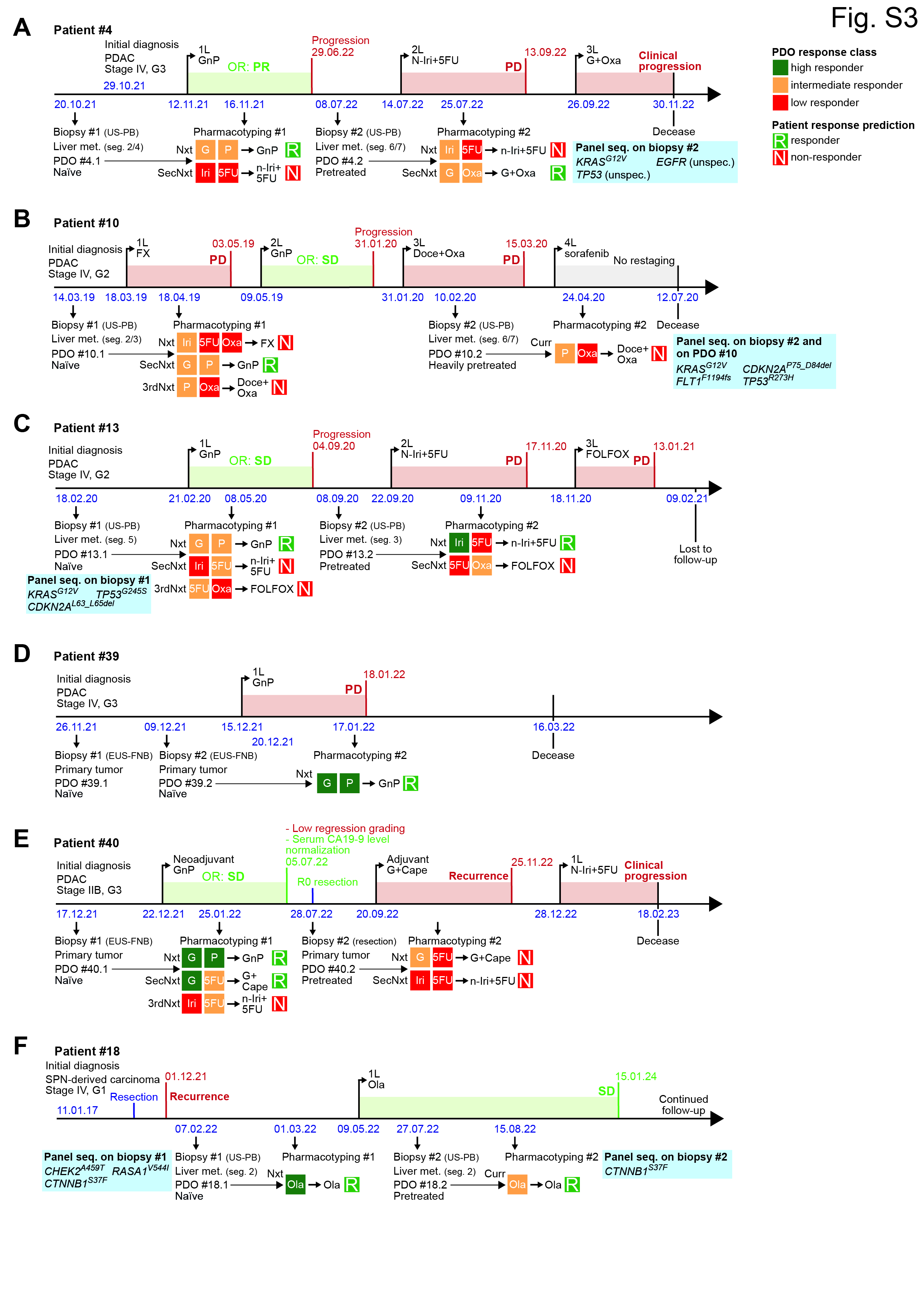

### Supplemental Figure S4

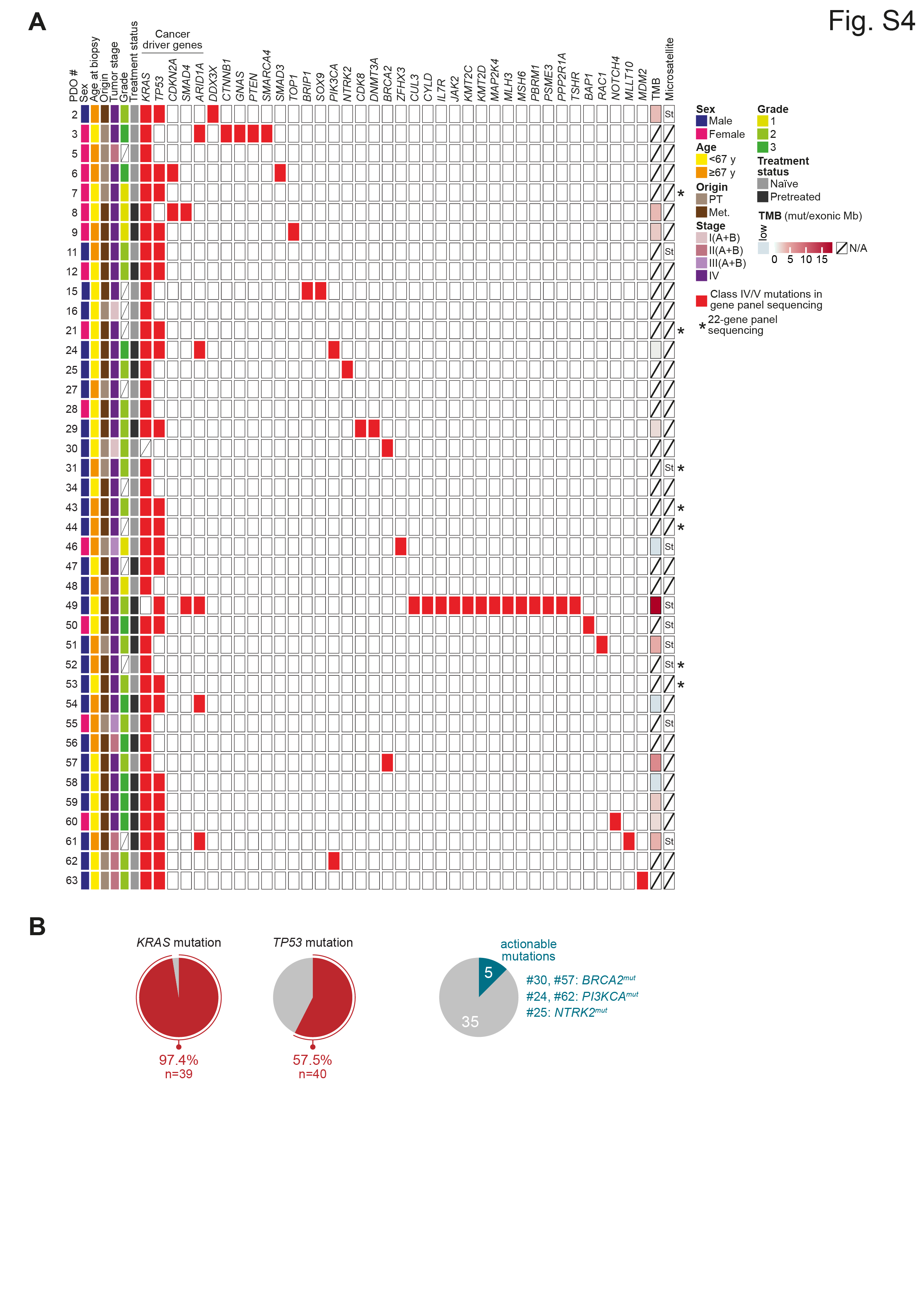

### Supplemental Figure S5

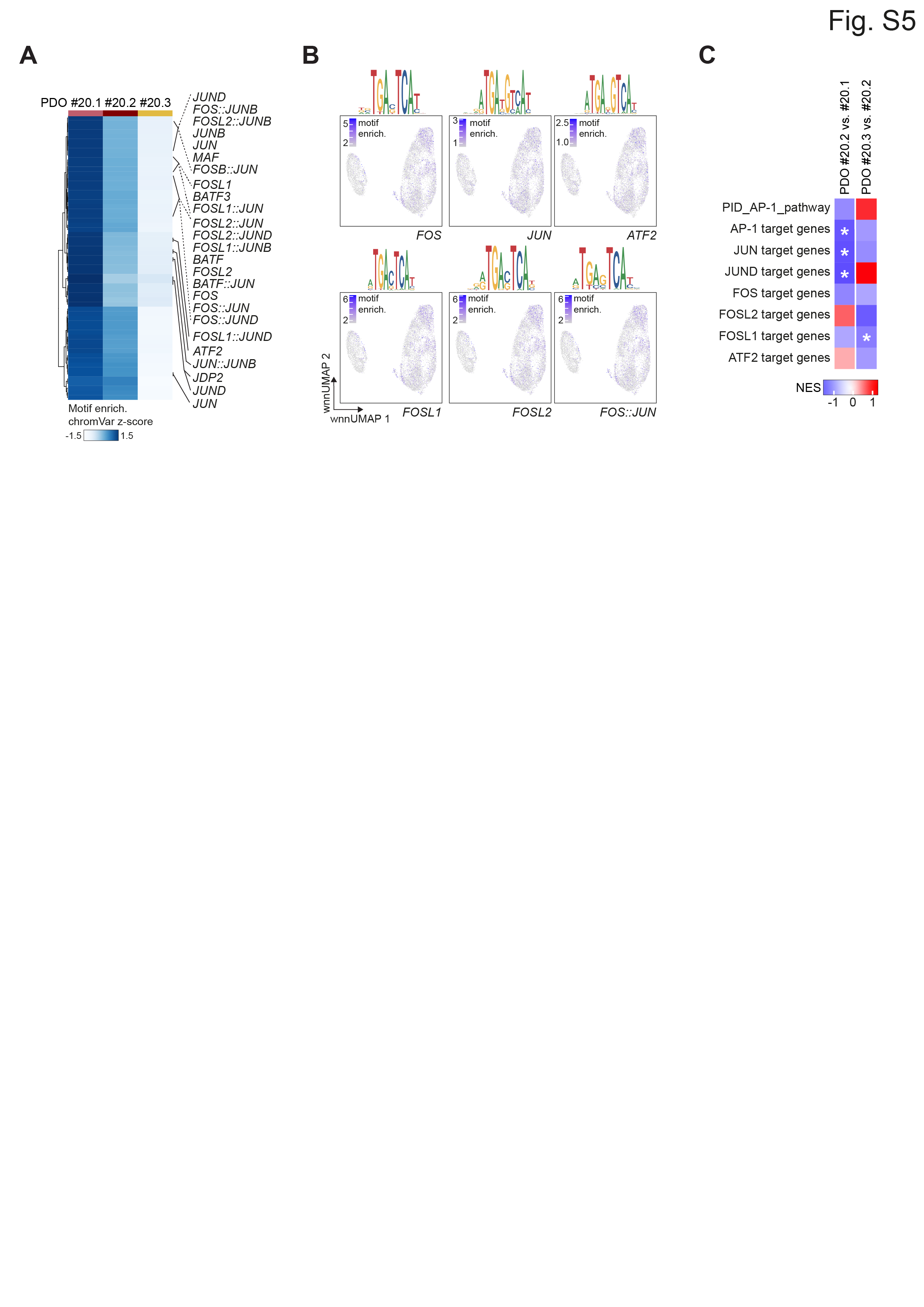

### Supplemental Figure S6

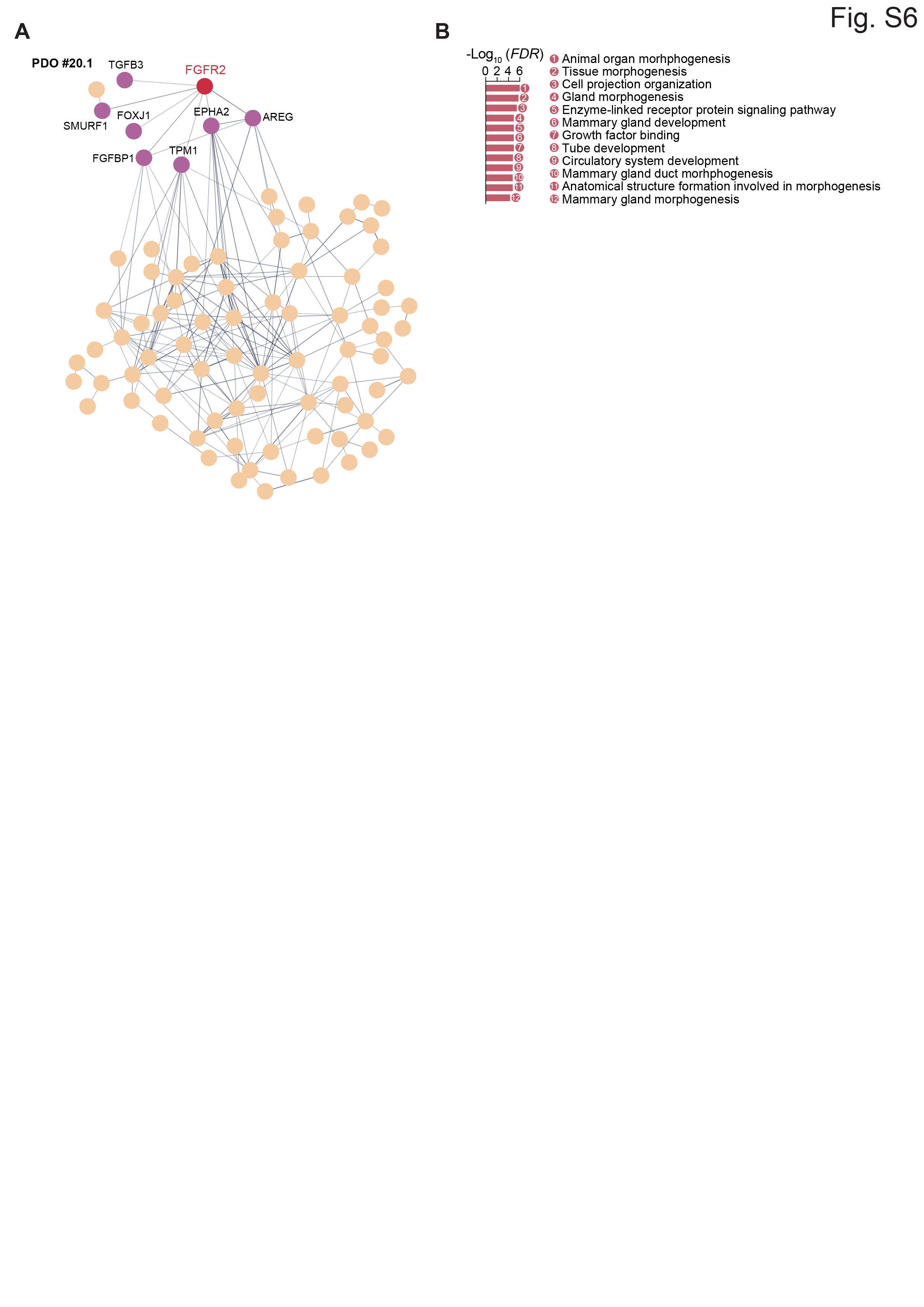
